# Origin of Ewing sarcoma by embryonic reprogramming of neural crest to mesoderm

**DOI:** 10.1101/2024.10.27.620438

**Authors:** Elena Vasileva, Claire Arata, Yongfeng Luo, Ruben Burgos, J. Gage Crump, James F. Amatruda

## Abstract

Ewing sarcoma is a malignant small round blue cell tumor of bones and soft tissues caused by chromosomal translocations that generate aberrant fusion oncogenes, most frequently EWSR1::FLI1. The cell of origin and mechanisms of EWSR1::FLI1-driven transformation have remained unresolved, largely due to lack of a representative animal model. By developing a zebrafish Ewing sarcoma model, we provide evidence for a neural crest origin of this cancer. Neural crest-derived cells uniquely tolerate expression of EWSR1::FLI1 and targeted expression of EWSR1::FLI1 in these cells generates Ewing sarcomas. Single-cell analysis of tumor initiation shows that EWSR1::FLI1 reprograms neural crest-derived cells to a mesoderm-like state, strikingly resulting in ectopic fins throughout the body. By profiling chromatin accessibility and genome-wide EWSR1::FLI1 binding, we find that the fusion oncogene hijacks developmental enhancers for neural crest to mesoderm reprogramming during cancer initiation. These findings show how a single mutation profoundly alters embryonic cell fate decisions to initiate a devastating childhood cancer.

## Introduction

Ewing sarcoma is a malignant bone and soft tissue tumor occurring in children, adolescents and young adults ^1^. Metastases are observed at the time of diagnosis in approximately 20-25% of patients, and less than one-third of patients with metastatic Ewing sarcoma will survive ^2^. Histologic examination of Ewing sarcomas typically reveals sheets of small round blue cells with a prominent nucleus and scant cytoplasm. The most important molecular feature of these tumors is the presence of reciprocal chromosomal translocations, which fuse members of the FET family of RNA-binding proteins (FUS, EWSR1, and TAF15) with various members of the ETS family of transcription factors. In 85% of cases, Ewing sarcoma cells carry the t(11;22)(q24;q12) translocation, resulting in the EWSR1::FLI1 fusion gene ^3,4^. The EWSR1::FLI1 oncofusion plays a crucial role in sarcoma development. In primary tumors, the oncofusion abnormally binds “promoter-like” and “enhancer-like” GGAA microsatellite repeats that are, for instance, present around loci of key Ewing sarcoma marker genes such as *NKX2-2*, *NR0B1*, and *CD99*. Additionally, EWSR1::FLI1 demonstrates similar affinity and specific binding to consensus single GGAA-containing ETS sites as the normal FLI1 protein ^5–8^. While EWSR1::FLI1 functions as a potent chromatin remodeler, disrupting the expression of numerous genes involved in cell-cycle control, cell migration, and proliferation, it remains unclear whether single GGAA and/or GGAA microsatellite repeats are necessary for EWSR1::FLI1-induced tumor initiation.

Ewing sarcoma was described in 1921 by James Ewing as an "endothelioma of the bone" (Ewing J, 1921). However, attempts to model Ewing sarcoma in mouse have not been successful ^9^, leading to theories that the human-specific GGAA microsatellite repeats may preclude development of a physiological animal model. In the absence of an animal model, definitive evidence for the cell of origin of Ewing sarcoma and the mechanisms for tumor initiation have remained elusive. Studies using primary tumors or in vitro cell culture have suggested that bone marrow mesenchyme and/or neural crest-derived cells (NCCs) may serve as potential cells of origin for Ewing sarcoma ^3^. Although expression of EWSR1::FLI1 is toxic in most cell types, both human bone marrow mesenchymal cells and neural crest-like cells can tolerate at least some degree of EWSR1::FLI1 expression in vitro ^10,11^. In vivo, both cell types have the capacity to differentiate into various connective and skeletal tissues, although mesenchymal potential of NCCs is normally restricted to the cranial region. It was shown that EWSR1::FLI1 can transform primary bone marrow–derived mesenchymal cells in vitro and generate tumors that display hallmarks of Ewing sarcoma in a mouse xenograft model ^12,13^. Moreover, transcriptomic profiles of different Ewing sarcoma cell lines with experimentally downregulated EWSR1::FLI1 expression converge toward that of mesenchymal cells with in vitro potential to give rise to adipocytes and osteoblasts ^14^. On the other hand, Ewing sarcoma cells express neuroectodermal markers, and overexpression of EWSR1::FLI1 in a rhabdomyosarcoma cell line upregulated genes in common with embryonic NCCs ^15^. However, as these previous studies largely examined the end state of Ewing sarcoma tumor-derived cells, it is difficult to extrapolate gene expression and differentiation potential to pinpoint the cell of origin for tumor initiation.

We previously characterized a mosaic genetic model of Ewing sarcoma via non-tissue-specific, Cre-inducible expression of human EWSR1::FLI1 in developing zebrafish ^16^. To more precisely control expression of EWSR1::FLI1 and to address the cell of origin, here we develop a stable transgenic fish line for Cre-inducible expression of EWSR1::FLI1 in NCCs. The neural crest is a transient, multipotent cell population that undergoes an epithelial to mesenchymal transition and extensive migration during early vertebrate development. In the cranial region, NCCs give rise to connective and skeletal structures, nerves, pigment cells, and other cell types. In contrast, trunk NCCs have a more restricted potential, generating sensory and sympathetic ganglia, adrenal chromaffin cells, and, via the dorsal pathway, melanocytes (Bronner & LeDouarin, 2012; Le Douarin & Dupin, 2003; Sommer, 2010). We find that expression of EWSR1::FLI1 in embryonic trunk NCCs results in loss of neuronal and glial markers and a concomitant gain of mesodermal expression, including the early mesoderm specifier *tbxta* (*T/Brachyury*) and mesenchymal genes *pdgfra*, *twist1a*, and *prrx1a*. Dramatic evidence of mesodermal reprogramming is seen in the fact that EWSR1::FLI1-expressing cells induce the formation of ectopic fins, some of which are subsequently replaced by Ewing sarcoma tumors. In addition, we show that EWSR1::FLI1 promotes neural crest to mesoderm reprogramming and tumor initiation by binding to ETS sites in developmental enhancers of mesodermal genes. Our work in a zebrafish model of Ewing sarcoma reveals that the EWSR1::FLI1 oncofusion hijacks normal developmental mesoderm enhancers in trunk NCCs to drive tumor initiation.

## Results

### NCCs selectively tolerate EWSR1::FLI1 expression in vivo

We previously used embryonic injection of Cre recombinase and Tol2 transposase mRNAs along with a ubiquitous ubi:loxP-DsRed-STOP-loxP-GFP-2A-EWSR1::FLI1 transposon expression plasmid (hereafter, ubi:RSG-2A-EF1) to drive mosaic expression of separate GFP and EWSR1::FLI1 proteins in zebrafish, resulting in small round blue cell tumors similar to human Ewing sarcoma in 30-40% of fish (Fig. 1A)^16^. To better understand how EWSR1::FLI1 might initiate tumors, we analyzed which embryonic cell populations tolerated the oncofusion protein. We first performed time-lapse imaging of ubi:RSG-2A-EF1-injected embryos versus ubi:loxP-DsRed-STOP-loxP-GFP-injected controls (ubi:RSG) from 5 hours post-fertilization (hpf) (approximately 50% epiboly stage) to 24 hpf (Fig. 1B, Fig. S1A,B, Movies 1, 2). In ubi:RSG controls, GFP expression was detected from 5 hpf onwards and was found throughout various tissues of the embryo in a mosaic fashion. In contrast, GFP-2A-EF1 expression resulted in high embryonic mortality ^16^. In surviving embryos, GFP-2A-EF1+ cells were largely restricted to dorsal and ventral regions of the trunk at 24 hpf. These data suggest the presence of cell populations in these regions that can selectively tolerate expression of the oncofusion.

**Figure 1.**
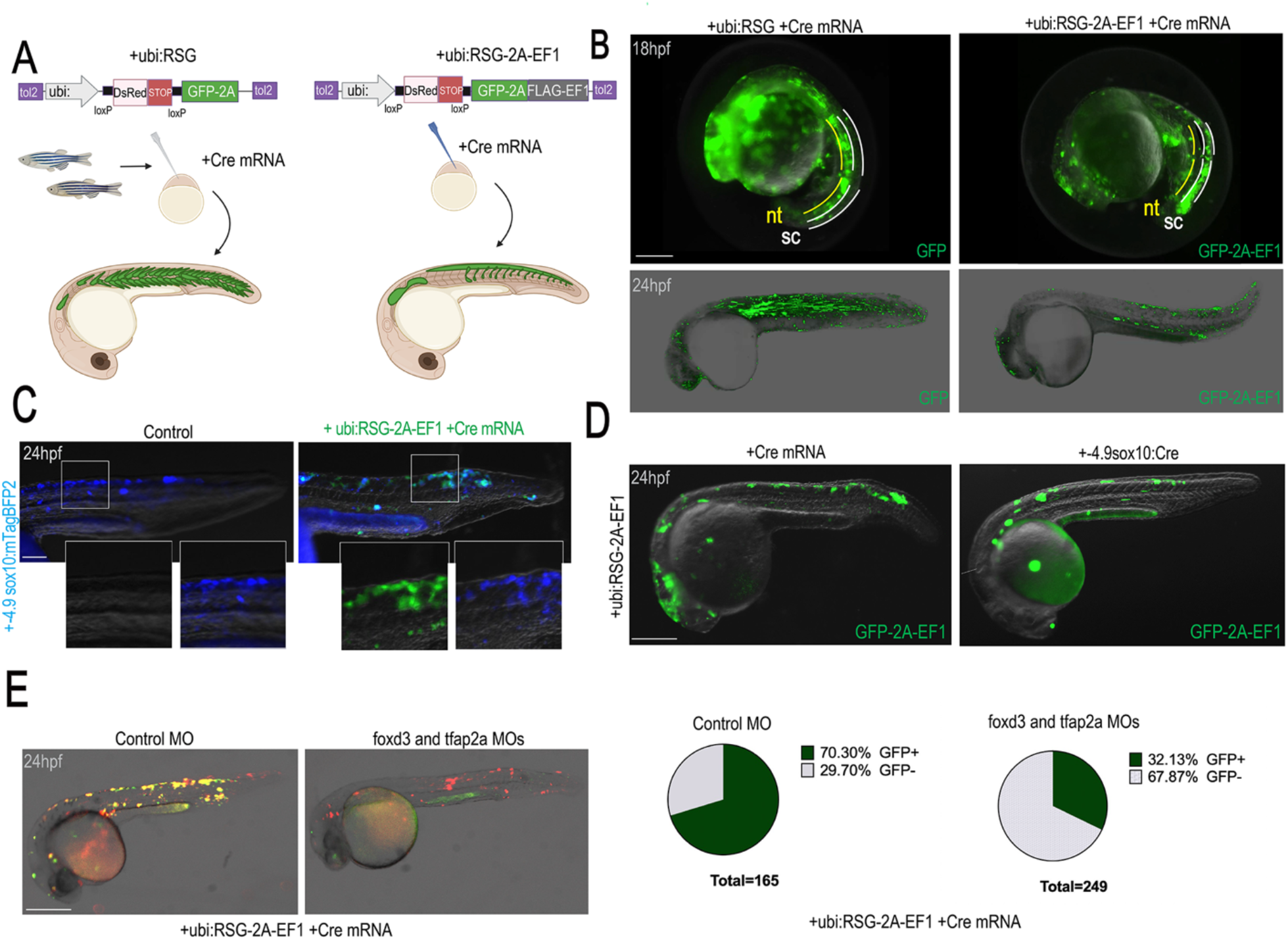
NCCs tolerate human EWSR1::FLI1 expression in vivo. **(A)** Overview of experimental method. The Tol2 transposon system was used to integrate constructs for Cre- inducible expression of GFP (control) and GFP-2A-EF1 into the wild-type WIK zebrafish genome by microinjecting into single-cell-stage embryos. Scale bars, 200 μm **(B)** Representative examples of serial imaging of fish expressing GFP (left) or GFP-2A-EF1 (right) at different stages of zebrafish development. sc: spinal cord; nt: notochord. Scale bars, 100 μm. **(C)** Representative images of 24 hpf embryos expressing -4.9sox10:mTagBFP2 alone (left) or co-expressing GFP- 2A-EF1 (right). Insets of individual channels show double-positive GFP-2A-EF1/mTagBFP2 cells. Scale bars, 100 μm. **(D)** Distribution of GFP-2a-EF1 positive cells using ubiquitous (ubi:RSG-2A-EF1+Cre mRNA) and NCC-specific (ubi:RSG-2A-EF1+-4.9sox10:Cre) mosaic models. Scale bars, 250 μm. **(E)** Embryos expressing GFP-2A-EF1 in the presence of control or *foxd3*/*tfap2a* morpholinos (MOs) which block development of NCCs. Embryos injected with *foxd3*/*tfap2a* MOs (N=249) show fewer GFP-2A-EF1+ cells (green) compared to embryos injected with control MO (N=165). The red channel shows unconverted ubi:RSG-2A-EF1-expressing cells unaffected by *foxd3*/*tfap2a* MOs. Scale bars, 250 μm.

As the location of GFP-2A-EF1+ cells in surviving embryos was reminiscent of trunk NCCs, we tested whether surviving cells had a NCC identity. Co-injection of the ubi:RSG-2A-EF1 cocktail with a construct consisting of NCC-restricted zebrafish -4.9 kb *sox10* promoter ^20^ driving TagBFP2 revealed double-positive cells (Fig. 1C, Fig. S2A, Movie 3). We also observed that a subset of pre- migratory and migratory NCCs co-expressed *sox10* RNA and GFP-2A-EF1 at 24 hpf (Fig. S2B). Further, co-injection of one-cell-stage embryos with the ubi:RSG-2A-EF1 plasmid and either Cre mRNA or the NCC-restricted -4.9sox10:Cre plasmid resulted in similar patterns of GFP-2A-EF1 expression (Fig. 1D). To confirm NCC identity of GFP-2A-EF1+ cells, we blocked NCC development by one-cell-stage embryo injection of morpholinos targeting *foxd3* and *tfap2a*, which replicate the NCC-less phenotype of *foxd3*; *tfap2a* genetic mutants ^21^. We validated NCC loss by injection into -28.5Sox10:Cre; actab2:loxP-BFP-STOP-loxP-DsRed (Sox10>DsRed) zebrafish embryos, in which NCCs are permanently labeled by DsRed ^22,23^ (Fig. S2C). Compared to co-injection of control morpholinos with the ubi:RSG-2A-EF1 cocktail, co-injection of *foxd3* and *tfap2a* morpholinos led to a decreased proportion of embryos expressing GFP-2A-EF1 (Fig. 1E). Taken together, these results indicate that NCCs selectively tolerate expression of EWSR1::FLI1.

### Ewing sarcoma formation by neural crest-specific expression of EWSR1::FLI1

To test whether expression of EWSR1::FLI1 in NCCs is sufficient to generate Ewing sarcomas, we generated -28.5Sox10:Cre; ubi:RSG-2A-EF1 fish to drive NCC-specific expression of GFP- 2A-EF1 (Sox10>GFP-2A-EF1) (Fig. 2A). As the zebrafish -4.9sox10 promoter has been shown to drive additional expression in cartilage and other tissues of mesodermal origin^24^, we used the murine -28.5Sox10 promoter that is more specific for NCCs in early zebrafish development ^22^ to drive Cre expression. In control Sox10>DsRed embryos at 72 hpf, DsRed expression was observed in NCC-derived Rohon-Beard (RB) and dorsal root ganglia (DRG) neurons and glial Schwann cells, as expected (Fig. 2B, left panel). In contrast, we observed GFP-positive cells not only in normal NCC positions but also as masses in the dorsal fin fold in Sox10>GFP-2A-EF1 embryos (Fig. 2B, right panel). Sequential live imaging from 72 hpf to 1 month revealed expansion of GFP+ fin fold masses (Fig. 2C). At 3-12 months of age, we observed small round blue cell tumors in Sox10>GFP-2A-EF1 fish that stained with antibodies specific for CD99, a cell surface glycoprotein that serves as a clinically useful marker for Ewing sarcoma ^25^ (Fig. 2D,E). Human Ewing sarcoma tumors also typically contain glycogen, and we confirmed the presence of glycogen in zebrafish tumors by Periodic acid-Schiff (PAS) staining ^25^. In addition, we confirmed continued expression of GFP-2A-EF1 in tumors by anti-GFP staining and nuclear localization of EWSR1::FLI1 by antibodies against human FLI1. Overall, these data show that expression of EWSR1::FLI1 in NCCs can lead to cell transformation in vivo and Ewing sarcoma formation.

**Figure 2.**
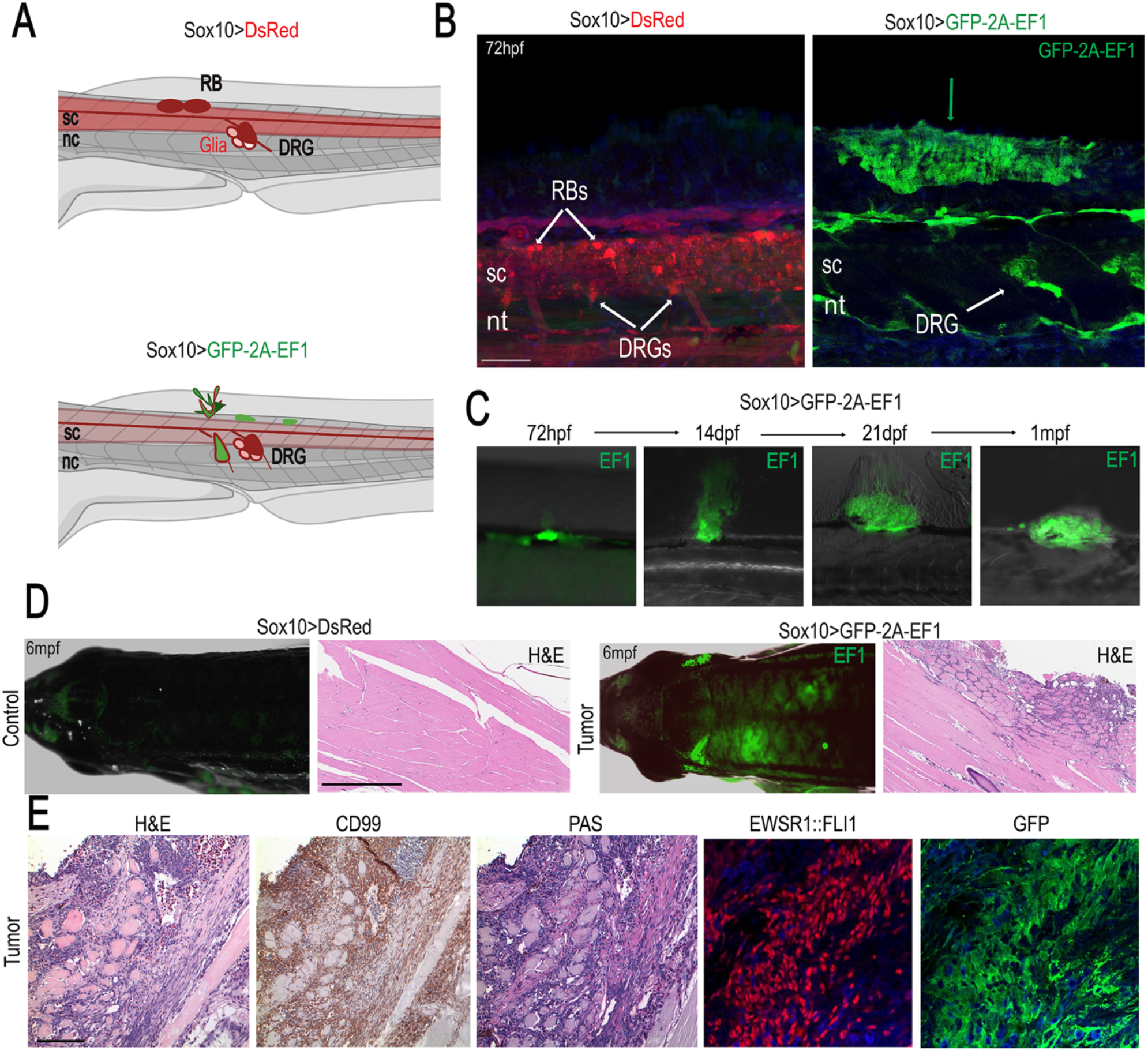
EWSR1::FLI1 expression in NCCs drives tumor development. **(A)** The - 28.5Sox10:cre; actab2:loxP-BFP-STOP-loxP-DsRed (Sox10>DsRed) transgenic line labels NCC derivatives, including Rohan-Beard (RB) and dorsal root ganglia (DRG) neurons and glia. Crossing this line to a stable ubi:loxP-DsRed-STOP-loxP-GFP-2A-EWSR1::FLI1 generates Sox10>GFP-2A-EF1embryos expressing EWSR1::FLI1 in NCCs. sc: spinal cord; nt: notochord **(B)** Expression of EWSR1::FLI1 in NCCs drives the formation of outgrowths on the median fin fold (right), which are not observed in control fish (left) at 72 hpf. EWSR1::FLI1 positive cells were also detected in the dorsal root ganglion (DRG). Scale bars, 50 μm. **(C)** Serial imaging of EWSR1::FLI1-expressing fish shows progression of GFP+ outgrowths from 72 hpf to 1 mpf. **(D)** Representative examples of control Sox10>DsRed, and a Sox10>GFP-2A-EF1 fish with tumors. Scale bars, 500 nm. **(E)** Characterization of NCC-derived tumors by staining for CD99, PAS, EWSR1::FLI1, and GFP. Scale bars, 100 μm.

### Trunk NCC-derived tumors exhibit mesenchymal features

In order to characterize NCC-derived tumors, we examined global gene expression and chromatin accessibility. To do so, we dissected a tumor from Sox10>GFP-2A-EF1 fish at 3 mpf and compared it to healthy tissue from the dorsal trunk of wild-type fish at the same age. We then performed combined single-nuclei RNA sequencing (snRNAseq) and single-nuclei assay for transposase-accessible chromatin sequencing (snATACseq) on dissociated cells using the 10x Genomics platform and Illumina sequencing (Fig. 3A). After filtering for quality, we obtained 2,884 cells with a median of 3,308 fragments and median 550 genes per cell for the tumor sample and 3,562 cells with a median of 4,058 fragments and 762 genes per cell for the control sample (Table S1). By analyzing tumor and control samples, we found that the datasets largely formed non-overlapping clusters composed of various cell types (Fig. 3B). The largest cluster in the tumor sample was composed of cells positive for EWSR1::FLI1 (which we term “tumor”). We also observed EWSR1::FLI1 expression in smaller populations of macrophages, immune cells, and periderm cells of the tumor sample, yet expression was absent in the control dataset (Fig. 3C). In both the main tumor cluster and a mesenchymal cluster from the control trunk, we observed expression of mesenchyme markers *pdgfra*, *twist1a*, and *prrx1a*. In contrast, the early mesoderm specifier *tbxta* was strongly expressed in tumor cells but absent in the control mesenchyme cluster (Fig. 3C). Reflecting selective expression of *tbxta* and other genes only in the tumor cluster, tumor and control mesenchymal cells formed distinct clusters (Fig. 3D). Consistently, gene ontology enrichment analysis of tumor versus control cells revealed terms related to early mesoderm development, such as ‘skeletal system development’, ‘fin development’, ‘notochord development’, and ‘somitogenesis’ (Fig. 3E). We also observed expression of orthologs of known human target genes of EWSR1::FLI1 in tumor but not control cells, including *cav1*, *fn1a*, *tnc*, *snx18a*, *fzd4*, *cdh11*, *igf1rb*, *fli1a*, *col21a1*, *SLCO5A1A*, and *dlg2* (Fig. S3) ^26,27^. These findings reveal that EWSR1::FLI1+ tumor cells originating from trunk NCCs express early mesodermal and mesenchymal genes, as well as a suite of genes associated with mature Ewing sarcomas in humans.

**Figure 3.**
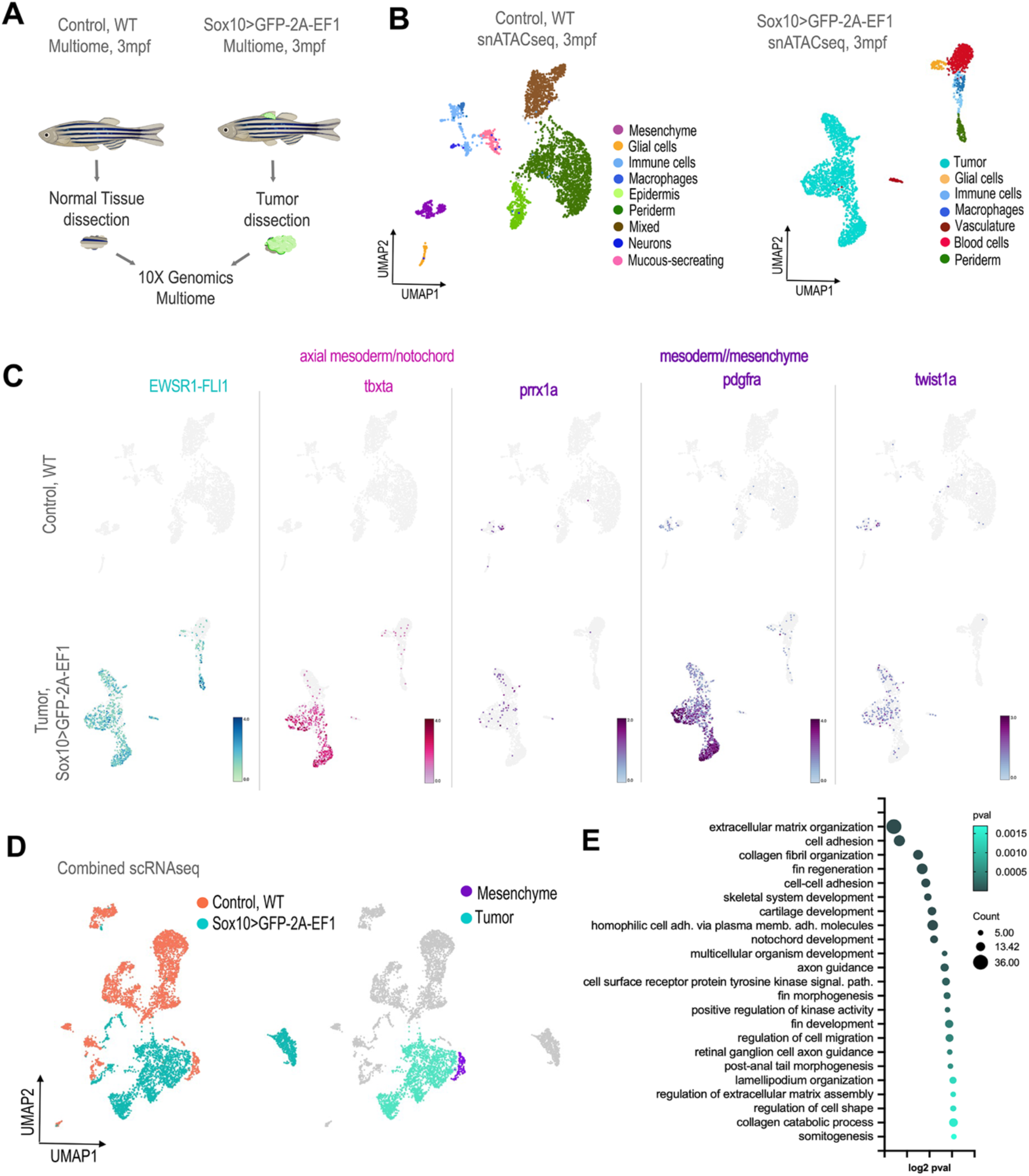
Mesenchymal features of NCC-derived adult tumors. **(A)** Schematic of sample processing of control tissue (wild type) and EWSR1::FLI1-expressing tumor tissue (Sox10>GFP- 2A-EF1) for combined snRNAseq and snATACseq (10x Genomics Multiome)**. (B)** UMAPs of control (4,058 cells) and tumor (3,308 cells) datasets. **(C)** Feature plots show the expression of mesodermal/mesenchymal markers in control and tumor datasets. **(D)** UMAPs of combined control and tumor datasets (left) and tumor and control mesenchyme clusters (right). **(E)** Gene Ontology enrichment analysis of genes differentially upregulated in tumor cells.

### EWSR1::FLI1 reprograms trunk NCCs to a mesoderm-like state

To characterize the earliest stages of NCC transformation mediated by EWSR1::FLI1, we next performed snRNAseq of Sox10>GFP-2A-EF1 versus Sox10>DsRed control cells at 7 dpf. GFP+ (EWSR1::FLI1-expressing) or DsRed+ (control) cells were isolated from dissected trunks by fluorescence-activated cell sorting (FACS), and snRNAseq sequencing was performed on the 10x Genomics Chromium platform followed by paired-end Illumina next-generation sequencing (Fig. 4A). After filtering for quality, we obtained 1,835 DsRed+ control and 388 GFP-2A-EF1+ cells, which largely separated into distinct clusters (Fig. 4B). Compared to control cells, EWSR1::FLI1- expressing cells had reduced expression of markers of neurons (*elavl4, isl2b*) and glia (*sox10, foxd3*), and higher expression of markers of embryonic mesenchyme (*pdgfra*, *twist1a*) and mesoderm (*tbx1, tbxta*) (Fig. 4C). Gene Ontology (GO) term enrichment analysis on differentially expressed genes (DEGs) in EWSR1::FLI1-expressing cells revealed a general downregulation of processes associated with neuronal development, including NCC-derived neurons (e.g., "enteric nervous system development") and neuronal function (e.g., “ion transport”, “chemical synaptic transmission”, “vesicle fusion”) (Fig. 4D).

**Figure 4.**
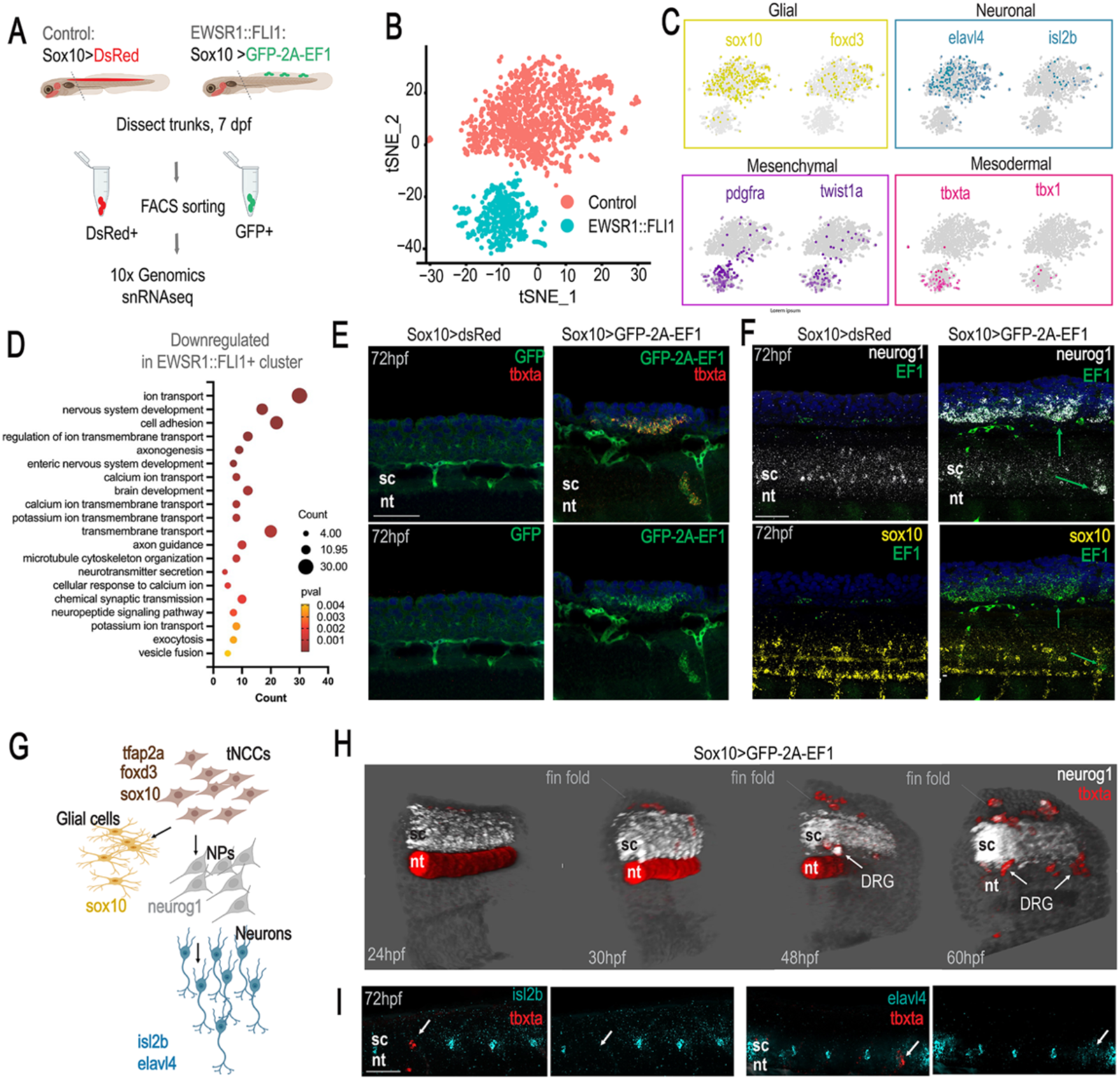
**EWSR1::FLI1 reprograms trunk NCCs to a mesoderm-like state and impairs neuronal development**. **(A)** Schematic of experimental design. **(B)** t-distributed stochastic neighbor embedding (t-SNE) plot of integrated Sox10>GFP-2A-EF1 (517 cells) and control Sox10>DsRed (1,854 cells) datasets. **(C)** Expression of glial (*sox10*, *foxd3*), neuronal (*elavl4*, *isl2b*), mesenchymal (*pdgfra*, *twist1a*), and mesodermal (*tbxta*, *tbx1*) markers in control and EWSR::FLI1+ cells. **(D)** Gene Ontology enrichment analysis of genes downregulated in the EWSR1::FLI1 cluster versus control cells. **(E)** GFP-2A-EF1 and *tbxta* expression in control Sox10>DsRed and Sox10>GFP-2A-EF1 fish. Scale bars, 50 μm. **(F)** Expression of markers of neuronal progenitor cells (*neurog1*) and glial cells (*sox10*) in control Sox10>DsRed and Sox10>GFP-2A-EF1 fish. Scale bars, 50 μm. **(G)** Schematic of trunk NCC differentiation into glial and neuronal lineages. **(H)** 3D reconstructions of *neurog1* (white) and *tbxta* (red) expression in larvae with Sox10>GFP-2A-EF1 driven outgrowths at 24-60 hpf. *tbxta*+ cells are observed in dorsal fin fold masses and DRGs. sc: spinal cord; nt: notochord. **(I)** In situ hybridization of Sox10>GFP-2A-EF1 fish for *tbxta* and neuronal markers *elavl4* and *isl2b*. Arrows show DRGs expressing *tbxta* but not *elavl4* or *isl2b*. Scale bars, 50 μm.

Tbxta/Brachyury plays an essential role during early gastrulation, where it directs posterior mesoderm and notochord formation ^28–30^. Consistent with snRNAseq results, RNA in situ hybridization revealed expression of *tbxta* in EWSR1::FLI1+ cells within the ectopic masses of the dorsal fin fold, as well as in normal DRG positions. We did not observe expression of *tbxta* in trunk NCCs or the dorsal fin fold of Sox10>DsRed controls at 72 hpf (Fig. 4E). 3D reconstructions from 24-60 hpf revealed that EWSR1::FLI1+/*tbxta*+ cells were adjacent to the spinal cord dorsally and laterally but were not observed within the spinal cord itself (Fig. 4H). In normal NCC development, *neurog1*-positive neural precursors give rise to mature neurons, with trunk NCC-derived DRG neurons adopting a glial fate in *neurog1* mutants ^31^ (Fig 4F,G). We therefore examined whether EWSR1::FLI1-expressing trunk NCCs that ectopically expressed *tbxta* retained any neuroglial properties from their NCC origin. EWSR1::FLI1+; *tbxta*+ cells in both DRGs and the dorsal fin fold masses expressed the neuronal progenitor marker *neurog1* but were negative for the glial marker *sox10* and more mature neuronal markers *isl2b* and *elavl4* (Fig. 4F,H,I). However, compared to the NCC-specific model, *tbxta+* cells in the mosaic model did not express *neurog1* at 72 hpf (Fig. S4D, Movie 4), potentially reflecting the earlier expression of EWSR1::FLI1 in the mosaic model relative to delayed EWSR1::FLI1 expression in trunk NCCs in the Sox10>GFP-2A- EF1 model (Fig. S4C). These findings indicate that EWSR1::FLI1 transforms trunk NCCs to a mesoderm-like state, with the exact timing of EWSR1::FLI1 expression determining whether the transformation occurs in uncommitted NCCs or those in a *neurog1*+ neuronal progenitor state.

### EWSR1::FLI1 opens ETS-containing putative enhancers during tumor initiation and progression

To understand how EWSR1::FLI1 induces mesodermal gene expression in trunk NCCs during early stages of tumorigenesis, we performed snATACseq to identify chromatin regions with altered accessibility following NCC-specific EWSR1::FLI1 expression. To do so, we FACS-isolated Sox10>GFP-2A-EF1 and control Sox10>DsRed cells at 7 dpf and performed 10x Genomics single-nuclei ATAC followed by Illumina sequencing (Fig. 5A). After filtering for quality, we obtained 1,922 cells with a median of 4,647 fragments per cell for the EWSR1::FLI1 sample and 1,854 cells with a median of 1,696 fragments per cell for control (Table S1). When visualized by UMAP, the EWSR1::FLI1-expressing and control populations largely separated into distinct clusters (Fig. 5A).

**Figure 5.**
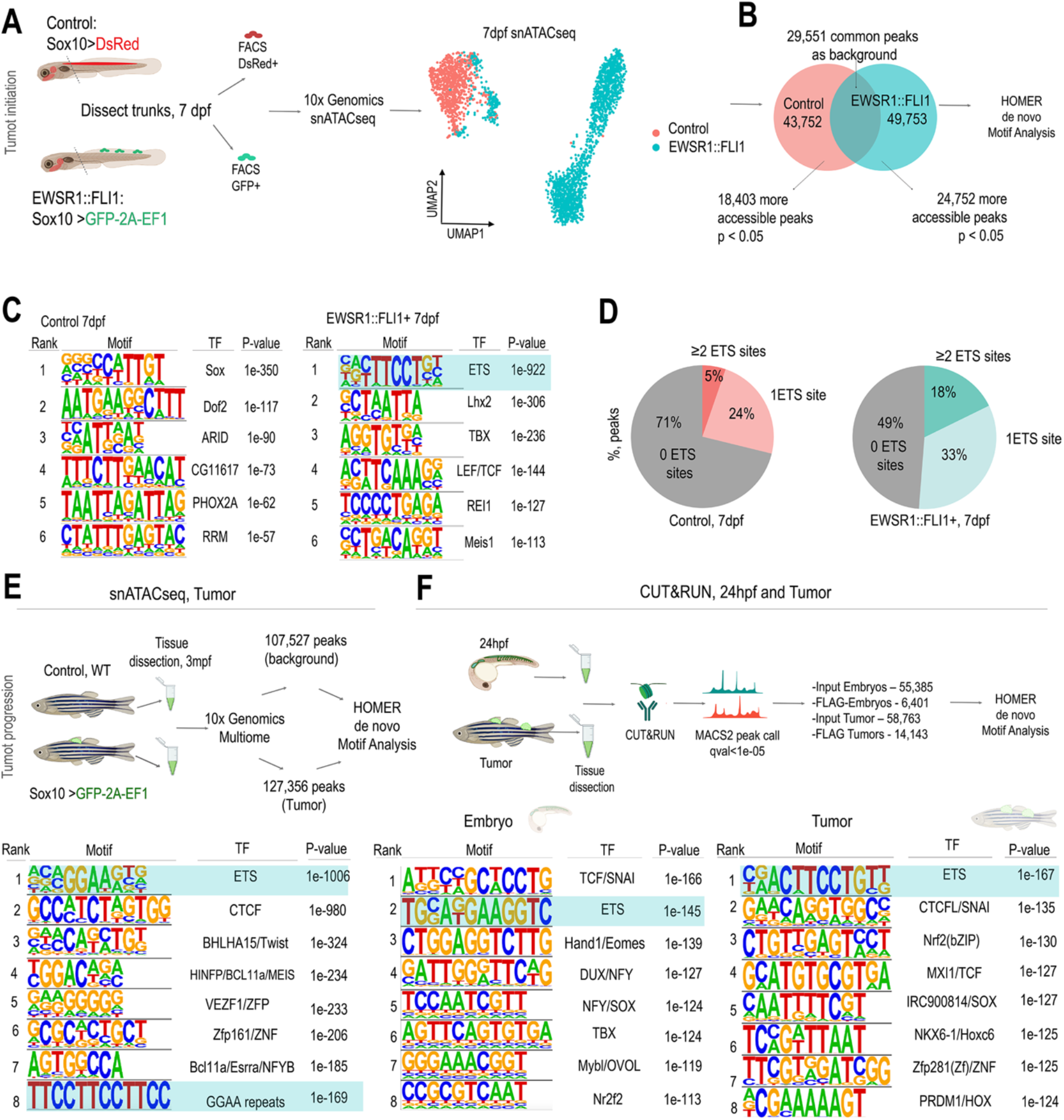
EWSR1-FLI1 binds ETS sites. **(A)** Schematic of sample processing of control (Sox10>DsRed) and EWSR1::FLI1 expressing (Sox10>GFP-2A-EF1) fish at 7 dpf for snATACseq. UMAP plot of integrated control (1,854 cells) and EWSR1::FLI1+ (1,922 cells) datasets. **(B)** More accessible peaks identified from Signac AccessiblePeaks for each dataset were used as input for HOMER de novo motif analysis, with common peaks used as background. **(C)** HOMER de novo motif analysis of more accessible peaks for control and Sox10>GFP-2A-EF1 datasets. **(D)** Quantification of the percentage of peaks with ETS sites (NRYTTCCTGH) in control and EWSR1::FLI1+ datasets, determined using HOMER annotatePeaks.pl. **(E)** HOMER de novo motif analysis of more accessible peaks in the tumor dataset (Sox10>GFP-2A-EF1) using peaks from the control dataset (normal tissue) as background. **(F)** HOMER de novo motif analysis of more accessible peaks from CUT&RUNseq using FLAG antibody on embryos mosaically expressing EWSR1::FLI1 at 24 hpf and on established tumors at adult stages, with Input datasets used as background.

We next used HOMER de novo motif enrichment analysis to identify transcription factor-binding motifs enriched in chromatin regions with increased accessibility in EWSR1::FLI1-expressing cells (49,753 total accessible peaks) versus control cells (43,752 total accessible peaks). We compared more accessible EWSR1::FLI1+ peaks (24,752) or more accessible control peaks (18,403) against a background of commonly accessible peaks (29,551) (Fig. 5B). The top-ranked motif in the control sample was SOX (p-value 1e-350), consistent with roles of Sox10 for NCC-derived melanocyte, glial, and neuronal development ^20,32^, and the fifth-ranked motif was PHOX2A, which specifies trunk NCC-derived sympathetic neurons ^33^ (Fig. 5C). In contrast, the top-ranked motif for EWSR1::FLI1+ cells was ETS (p-value 1e-922). More accessible chromatin regions in EWSR1::FLI1+ cells were also enriched for motifs for TBX (3^rd^, p-value 1e-236) and the Wnt effector LEF/TCF (4^th^, p-value 1e-144), consistent with roles of Tbx proteins and Wnt signaling in mesoderm development ^29,34^. In addition, 51% of total accessible peaks (common peaks removed) had one or more ETS sites in EWSR1::FLI1+ cells, while only 29% of regions had ETS sites in controls (Fig. 5D). Although ETS sites contain GGAA as part of their core motif, GGAA repeats were not uncovered as a significant motif in EWSR1::FLI1+ cells at tumor initiation stages.

To understand whether this pattern of chromatin accessibility is maintained in tumors, we performed HOMER de novo motif enrichment analysis to identify transcription factor binding motifs enriched in open chromatin of tumors (127,356 accessible peaks) versus control trunk tissues (107,527 peaks) in our multiome datasets. (Fig. 5E). The top-ranked motif for tumor cells was ETS (p-value 1e-1006), a near perfect match for the ETS motif in open chromatin of 7 dpf EWSR1::FLI1-expressing cells (Fig. 5C). We also uncovered a GGAA repeat as the 8^th^ motif (p-value 1e-169), in contrast to open chromatin of 7 dpf EWSR1::FLI1-expressing cells that was not enriched for GGAA repeat motifs. These findings show that EWSR1::FLI1 shifts from promoting accessibility of chromatin containing single GGAA-containing ETS sites at tumor initiation stages to chromatin containing both ETS and GGAA repeat sequences in mature tumors.

To determine which regions of accessible chromatin are directly bound by the oncofusion protein, we performed Cleavage Under Targets & Release Using Nuclease sequencing (CUT&RUNseq) during tumor initiation (24 hpf, 2 technical replicates) and maintenance (2-8 mpf, 2 pooled tumors). To capture sufficient cell numbers, we used the mosaic model (Fig. 5F). As the transgenic EWSR1::FLI1 protein contains a FLAG epitope, we performed CUT&RUNseq using an anti-FLAG antibody and performed HOMER de novo motif enrichment analysis using input peaks as background (Fig. 5F). At the embryonic timepoint (24 hpf), we uncovered ETS as the 2nd ranked motif and TBX as the 6th motif, in concordance with these motifs in the 7 dpf EWSR1::FLI1+ snATACseq dataset (Fig. 5C). For the tumor CUT&RUNseq dataset, we uncovered ETS as the top ranked motif and CTCF as the 2nd, in agreement with the tumor snATACseq dataset. These findings confirm our snATACseq results that EWSR1::FLI1 largely binds single GGAA-containing ETS sites during tumor initiation, potentially in combination with TBX and other developmental factors.

### EWSR1::FLI1 directly binds ETS sites of mesodermal developmental enhancers during tumor initiation and growth

Upregulation of mesodermal and mesenchymal gene expression in pre-tumor and tumor cells led us to investigate how the mesodermal program is regulated during tumor initiation and formation. To determine if the genomic regions of mesodermal (*tbxta*) and mesenchymal genes (*pdgfra*, *twist1a*, *prrx1a*) contain putative developmental enhancers bound by the oncofusion protein, we examined in more detail their genomic loci. We compared CUT&RUNseq and snATACseq datasets and complemented our data with publicly available ATACseq datasets from lateral plate mesoderm at 12 hpf and tail bud at 24 hpf ^35^ (Fig. 6). We observed regions of EWSR1::FLI1 binding within 50-100 kb of the transcription start sites (TSS) of the *tbxta*, *prrx1a*, *pdgfra*, and *twist1a* genes. The loci of all these genes were generally more accessible in EWSR1::FLI1- expressing versus control tissues. We uncovered two major types of accessible regions in EWSR1::FLI1-expressing cells that corresponded to accessible regions in the lateral plate mesoderm and/or tail bud of normal embryos. One type was directly bound and made accessible by EWSR1::FLI1, based on selective accessibility in EWSR1::FLI1-expressing versus control snATACseq datasets and direct binding by EWSR1::FLI1 in the CUT&RUNseq dataset. These putative direct developmental enhancers included at least 5 regions for the *tbxta* gene, two of which are known posterior mesoderm enhancers (Hox element 1 (HE1) and Hox element 2 (HE2)) ^36^. We also identified at least one direct developmental enhancer each for *pdgfra*, *twist1a*, and *prrx1a* genes. These putative enhancers varied in their EWSR1::FLI1 binding and chromatin accessibility at early and late stages, suggesting that they may open at different times of tumor initiation and stay accessible for different periods of time. We also observed putative indirect developmental enhancers that gained accessibility upon EWSR1::FLI1 expression but were not bound by EWSR1::FLI1 in CUT&RUNseq. All putative direct developmental enhancers contained predicted ETS binding sites. In addition, we observed EWSR1::FLI1 binding to the promoters of *tbxta*, *twist1a* and *prrx1a*, which also contained predicted ETS sites. Lastly, we observed that some of the regions of strongest EWSR1::FLI binding corresponded to regions that lacked chromatin accessibility in tumor, control, and normal developmental ATACseq datasets, suggesting that they are not active enhancers. Two of these regions contained GGAA repeat sequences, with one of these in the *prrx1a* locus also containing 31 predicted ETS sites. Thus, while EWSR1::FLI1 appears to also bind GGAA repeat sequences in zebrafish, our findings support EWSR1::FLI driving tumor initiation in trunk NCCs by binding to normal ETS-containing developmental mesoderm enhancers.

**Figure 6.**
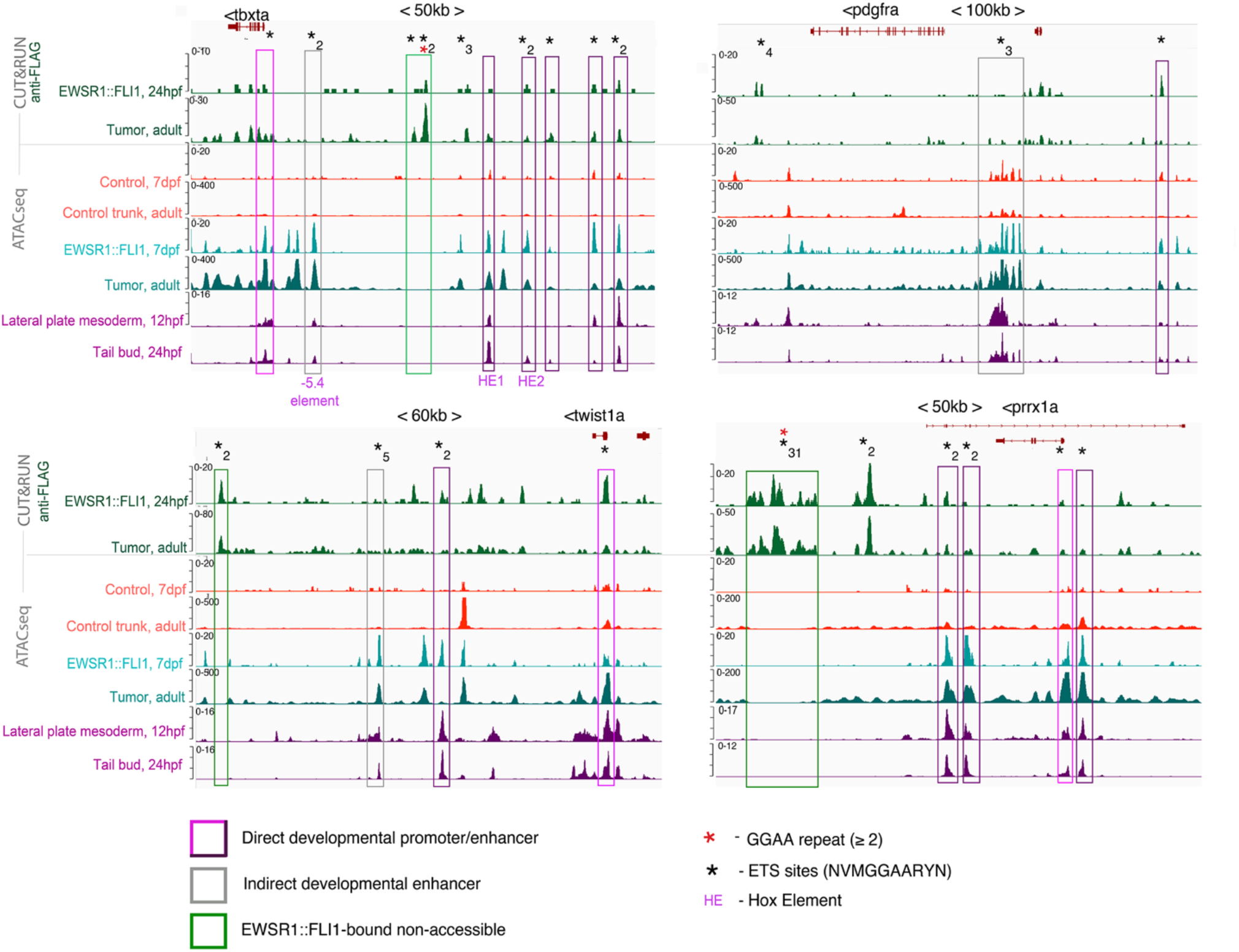
EWSR1::FLI1 directly binds developmental mesoderm enhancers. Integrated coverage plots for anti-FLAG CUT&RUNseq (EWSR1::FLI1+ at 24 hpf and in adult tumors) and snATACseq (control and EWSR1:: FLI1+ at 7 dpf, normal tissue and EWSR1::FLI1+ tumor at adult stages, and lateral plate mesoderm and tail bud downloaded from GEO Series GSE243256). Genomic loci for *tbxta*, *pdgfra*, *twist1a*, and *prrx1a* are shown. Directly bound developmental promoters/enhancers (pink/magenta), indirectly regulated developmental enhancers (gray), and regions bound by EWSR1::FLI1 but lacking chromatin accessibility in any dataset (green) are boxed. Black asterisks denote ETS sites (number indicated per region) and red asterisks mark GGAA repeats. Known enhancers -5.4 element, HE1, and HE2 are shown for *tbxta*.

### Transformed EWSR1::FLI1+ trunk NCCs can induce ectopic fins

The transcriptional reprogramming of trunk NCCs to a mesoderm-like state by EWSR1::FLI1 prompted us to investigate whether reprogrammed cells exhibited functional features of embryonic mesoderm, including the ability to induce fins. A time-course of *tbxta* expression from 24 hpf to 14 dpf revealed *tbxta*-expressing cells located adjacent to the spinal cord at 24 hpf and *tbxta*- expressing ectopic fin fold masses that grew in size in EWRS1::FLI1-expressing fish from 72 hpf to14 dpf but were absent in controls (Fig. 7A, Movie 5). We also observed expression of transgenic GFP-2A-EWRS1::FLI1 in the ectopic fin masses in both NCC-specific and mosaic models at 72 hpf (Fig. 2C, S4A,B, Movie 5), which grew in size by 7 dpf and had a clear fin morphology by 21 dpf (Fig. 7B). Whereas ectopic outgrowths were seen in the dorsal and ventral fin folds in the mosaic model, they were largely restricted to the dorsal fin fold in the NCC-specific model (Fig. S4C). Ectopic fins were also in general larger and more frequent in the mosaic versus the NCC-specific model, which correlated with lower adult viability in the mosaic model (Fig. 7C). Although EWSR1::FLI1-expressing cells could induce ectopic fins, they primarily remained at the bases of fins (Fig. 7B). In many cases, ectopic fins regressed later in development, coincident with contribution of EWSR1::FLI1-positive cells to small round blue cell tumors (Fig. S5A). In some cases, ectopic fins were maintained until the young adult stage (2 months post-fertilization), correlating with decreased GFP-2A-EWRS1::FLI1 expression (Fig. S5B). These persistent ectopic fins were patterned normally into segmented fin rays visible by Alizarin Red staining (Fig. S5C) and with the ability to regenerate following amputation (Fig. S5D). A subset of fish sorted for EWSR1::FLI1-positive outgrowths at embryonic stages also displayed a reduction or absence of normal fin structures at adult stages (Fig. S5B). These findings show that EWSR1::FLI1- transformed NCCs can function similarly to normal mesoderm to induce fins, although the persistent undifferentiated state of EWSR1::FLI1-positive cells at the bases of fins leads to tumor formation.

**Figure 7.**
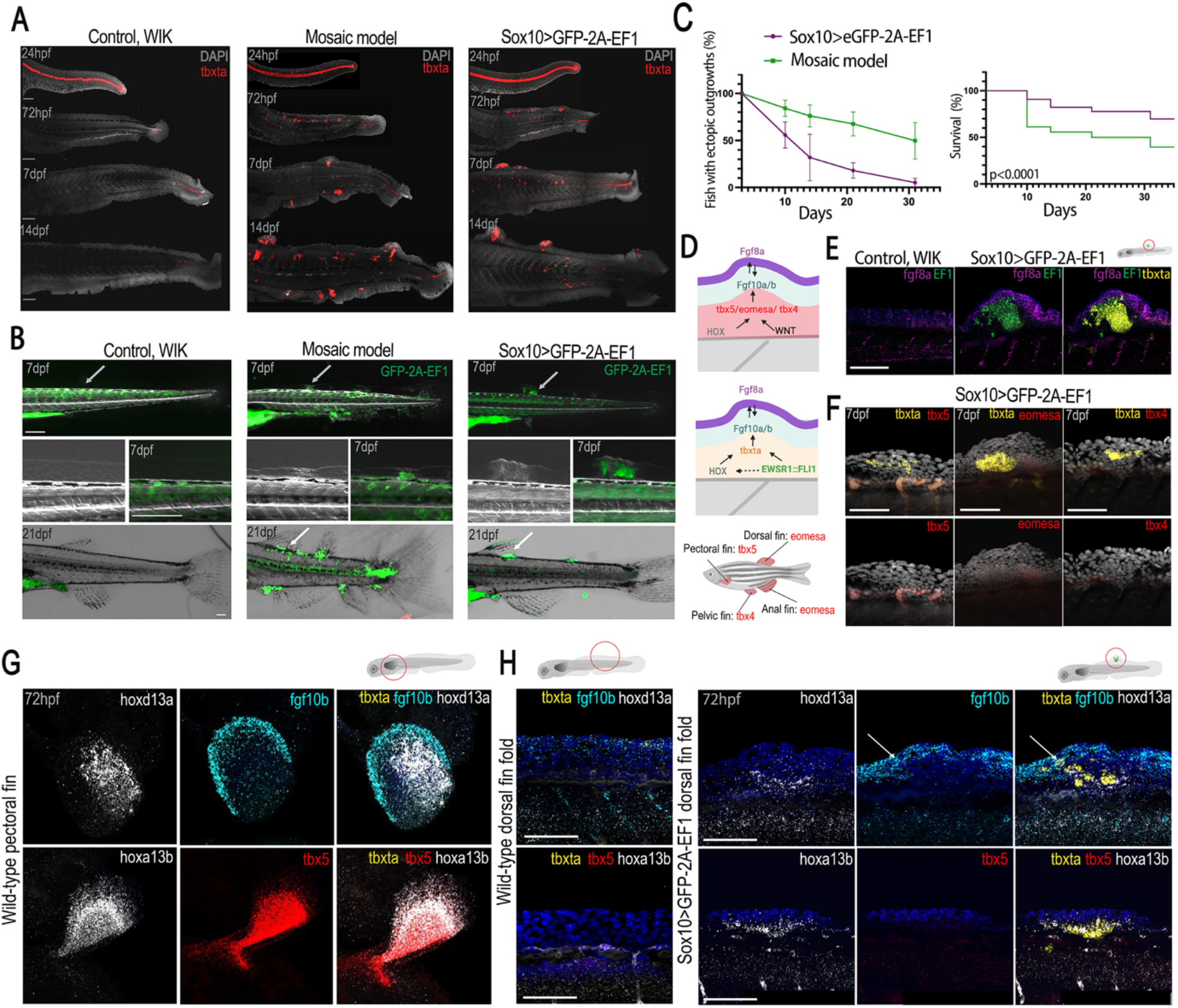
EWSR1::FLI1 activates an ectopic fin developmental program. **(A**) ISH for *tbxta* from 24 hpf to 14 dpf in non-transgenic controls, fish with mosaic expression of GFP-2A-EF1, and Sox10>GFP-2A-EF1 fish. *tbxta* is expressed in the notochord and tail mesoderm in control embryos. Ectopic expression of *tbxta* in trunk NCC migration domains and fin folds is observed in mosaic and Sox10>GFP-2A-EF1 models. Scale bars, 100 μm. **(B**) Expression of EWSR::FLI1 at 7 and 21 dpf in mosaic (ubi:GFP-2A-EF1) or NCC-specific (Sox10>GFP-2A-EF1) fish. Wildtype fish were used as control; green signal in wild types indicates autofluorescence. GFP- 2A-EF1-positive cell masses can be observed in the fin folds at 7 dpf, and at the bases of the ectopic fins at 21 dpf in both models. Arrows denote fin fold masses that developed into ectopic fins upon serial imaging of the same animals. Scale bars, 200 μm. (**C)** Percentage of fish with ectopic outgrowths (top) and the overall survival rate of fish (bottom) in mosaic (ubi:GFP-2A-EF1 + Cre mRNA, N=106, green) and NCC-specific (Sox10>GFP-2A-EF1, N=238, purple) models. (**D)** Schematic shows signaling pathways involved in fin development, including WNT, HOX factors, T-box transcription factors, Fgf8a, and Fgf10a/b. Lower schematic shows normal expression of T-box transcription factors in different fins. **(E)** Expression of *fgf8a*, *EF1*, and *tbxta* in dorsal fin fold of control and Sox10>GFP-2A-EF1 larvae. Scale bars, 100 μm **(F)** *tbxta* but not *tbx5*, *eomesa*, or *tbx4* is expressed in Sox10>GFP-2A-EF1-induced outgrowths. **(G)** Expression of *hoxd13a, fgf10b,* and *tbx5*, but not *tbxta*, in a pectoral fin of a control embryo at 72 hpf. Scale bars, 100 μm. (**H)** Expression of *hoxd13a, hoxa13b, fgf10b, tbx5*, and *tbxta* in the dorsal median fin fold of control and Sox10>GFP-2A-EF1 larva at 72 hpf. Arrow denotes cells co-expressing *tbxta* and *fgf10b*. Scale bars, 100 μm

To understand how EWSR1::FLI1-transformed trunk NCCs induced ectopic fins, we examined the expression of genes important for normal fin development. The apical ectodermal ridge (AER) promotes fin outgrowth through a WNT and FGF signaling cascade and expression of posterior Hox genes (e.g., *hoxa13b*, *hoxd13a*) and Tbx genes (*tbx4*: pelvic fin, *tbx5*: pectoral fin, *eomesa*: dorsal and anal fins) ^37–41^ (Fig. 7D). Compared to inter-fin regions of the control dorsal fin fold that lacked *fgf8a* expression, ectopic fin buds in Sox10>GFP-2A-EF1 and mosaic models displayed AER-like expression of *fgf8a* in the epithelium, which did not overlap with GFP-2A- EWSR1::FLI1 expression in the underlying mesenchyme (Fig. 7E, Fig. S6A,C). Whereas EWSR1::FLI1-driven fin buds did not express Tbx family members normally expressed in developing fins (*tbx4*, *tbx5*, *eomesa*), they did express *tbxta* at 3 and 7 dpf, as well as *hoxa13b and hoxd13a* (Fig. 7E,F,H; Fig. S6A,B). We also observed mesenchymal expression of *fgf10a* and *fgf10b* in ectopic fin buds, although only a subset of *fgf10a* and *fgf10b* expression overlapped with EWSR1::FLI1 (Fig. 7H, Fig. S6D). Analysis of CUT&RUNseq and snATACseq datasets indicate that EWSR1::FLI1 directly binds the promoter of the *fgf10a* gene (Fig. S6E). Thus, EWSR1::FLI1 expression in trunk NCCs induces ectopic fin formation through largely the same molecular pathways as normal fin development, with the exception that *tbxta* substitutes for typical Tbx genes.

## Discussion

The cell of origin of Ewing sarcoma has been a subject of longstanding debate. By developing a tissue-specific zebrafish model of Ewing sarcoma, we demonstrate that trunk NCCs uniquely tolerate the EWSR1::FLI1 oncofusion protein and can develop into tumors upon expression of the oncofusion. Moreover, we reveal that EWSR1::FLI1 reprograms trunk NCCs into an undifferentiated mesoderm-like state that is maintained into adulthood, with these cells inducing ectopic fins and eventually forming tumors with key hallmarks of human Ewing sarcoma. Mechanistically, we find that the oncofusion induces mesodermal reprogramming of trunk NCCs by binding ETS sites within developmental mesoderm enhancers. Our work thus reveals NCCs as a cell of origin for Ewing sarcoma and suggests that the oncofusion drives transformation by hijacking a normal mesodermal developmental program.

Our finding that NCCs can selectively tolerate EWSR1::FLI1 expression is consistent with reports that human NCC-like cells can tolerate the oncofusion in vitro ^11^. Recent reports have suggested that during their early development NCCs either retain or reacquire a pluripotency-like signature ^42,43^, which may explain their ability to form mesenchymal and neuroglial derivatives characteristic of different germ layers. In addition, when grafted to a cranial position, trunk NCCs can be reprogrammed to a mesenchymal state ^44^. One possibility then is that this inherent plasticity of trunk NCCs allows them to be transformed by EWSR1::FLI1 without cell death. A NCC origin may also help explain why human Ewing sarcomas often exhibit both neuroectodermal and mesenchymal gene expression and characteristics ^1,14,45,47^.

One of the advantages of our model is the ability to examine the earliest molecular events leading to EWSR1::FLI1-mediated transformation and tumor initiation, which are not accessible in human Ewing sarcomas. Previous work in primary tumors and cell culture models of Ewing sarcoma formation had suggested that EWSR1::FLI1 activates gene expression through binding to human-specific GGAA microsatellite repeats ^5^. While EWSR1::FLI1 had also been shown, similar to normal FLI1, to bind to consensus ETS sites that contain GGAA as part of its motif, such binding appeared to be largely repressive ^5–8,46^. By studying Ewing sarcoma in zebrafish, we found that EWSR1::FLI1 acts differently during the earliest stages of tumor initiation. In particular, we revealed that EWSR1::FLI1 binds ETS sites in normal developmental enhancers for mesoderm and mesenchymal genes such as *tbxta*, *pdgfra*, *twist1a*, and *prrx1a*, resulting in increased chromatin accessibility and activity of these enhancers in trunk NCCs. Although we did also detect strong EWSR1::FLI1 binding to GGAA repeats, these regions did not display chromatin accessibility in either transformed NCCs or normal mesodermal cell populations, suggesting they are not active enhancers. Our findings therefore suggest that EWSR1::FLI1 functions during tumor initiation to hijack normal ETS-containing mesodermal enhancers. As opposed to GGAA repeat sequences that are not conserved from human to mouse or zebrafish, the role of FGF signaling and its downstream ETS-containing transcription factors in mesoderm formation is deeply conserved across vertebrates ^48,49^. The hijacking of ETS-containing mesodermal enhancers by EWSR1::FLI1 in tumor initiation, as opposed to binding to human-specific GGAA repeat sequences, would explain why Ewing sarcoma can be effectively modeled in zebrafish.

The ability of trunk NCC-derived EWSR1::FLI1-expressing cells to induce ectopic fins is a dramatic manifestation of their transformation to a mesoderm-like identity. Normally, trunk NCCs do not contribute to mesenchymal derivatives ^17^, with zebrafish and medaka fin mesenchyme shown to have an exclusively mesodermal origin ^50,51^. In both mosaic and NCC-specific EWSR1::FLI1 misexpression models, we found that the oncofusion cell-autonomously induces expression of *hoxa13b* and *hoxd13a*, essential regulators of pectoral fin formation in zebrafish ^52,53^. It is known that HOX genes are dysregulated in human Ewing sarcoma ^54^, with upregulation of *HOXD13* playing a role in regulating the mesenchymal state of Ewing sarcoma tumor cells ^55^. We also found that misexpression of EWSR1::FLI1 in zebrafish directly induced expression of *tbxta (brachyury)* in ectopic fin buds through normal developmental enhancers. As Tbx genes implicated in normal fin development (*tbx4*, *tbx5*, *eomesa*) were not upregulated by EWSR1::FLI1, it is possible that Tbxta substitutes for these related family members in inducing ectopic fins ^56^. In addition, zebrafish tumors maintain expression of *tbxta*, and expression of *TBXT* is observed in over half of Ewing sarcomas ^57^, suggesting that human tumor cells may also retain characteristics of early mesoderm.

Both EWSR1::FLI1-expressing and neighboring non-expressing mesenchymal cells express *fgf10a* and *fgf10b* in ectopic find buds, and EWSR1::FLI1 directly binds promoter regions of *fgf10a* analogous to those in tetrapod *Fgf10* bound by EWSR1 and FLI1 during normal limb formation ^58^. Since an Fgf8-Fgf10 positive feedback loop is necessary and sufficient for limb formation ^59^, one possibility is that EWSR1::FLI1 initiates mesenchymal *fgf10a* expression that induces epithelial *fgf8a* expression, which then signals back to non-EWSR1::FLI1-expressing mesenchyme to further induce *fgf10a* and *fgf10b*. The growth of human Ewing sarcomas is heavily influenced by FGF signaling, both in tumor cells ^60,61^ and the bone microenvironment ^62^. Thus, by driving elements of a limb developmental program, EWSR1::FLI1 may create a local signaling environment that favors tumor growth. The association with and ability of EWSR1::FLI1- transformed cells to induce ectopic fins may also help explain why Ewing sarcoma in humans often occurs in bones and soft tissue of the limbs.

## Limitations of the study

EWSR1::FLI1-mediated transformation of NCCs was evident as early as 24 hpf and led to tumor formation during the first 2-6 months of life. This is consistent with Ewing sarcoma having an early embryonic origin, resulting in cancer in children and young adults with the peak of incidence between 15 and 20 years. However, we did not test whether induction of EWSR1::FLI1 in other cell types or NCC-lineage cells at later timepoints could also induce Ewing sarcoma. While our findings strongly support NCCs being a cell of origin for Ewing sarcoma, we cannot rule out that other cell types could also be a source of Ewing sarcoma. For example, mosaic embryo-wide EWSR1::FLI1 misexpression resulted in ectopic ventral fins that were not observed in the NCC-specific expression model. It is possible that a deeper investigation of human tumors could reveal the existence of multiple subclasses of Ewing sarcomas, sharing broadly similar histopathologic and molecular features, but with distinct developmental origins. Whereas EWSR1::FLI1- transformed NCCs induced ectopic fins in zebrafish, extra limbs or digits are not a reported feature of human Ewing sarcoma. Unlike in the genetic zebrafish model where the oncofusion is widely expressed in embryonic NCCs, in humans the translocation event may occur in only a small subset of NCCs insufficient to induce ectopic limbs. While our genomics studies identified several direct targets of EWSR1::FLI in NCCs, future studies will also be needed to determine which of the many targets of EWSR1::FLI1 are critical for tumor initiation and growth.

## Materials and Methods Zebrafish husbandry

*Danio rerio* were maintained in adherence to established industry standards within an AALAAC-accredited facility. WIK wild-type fish were sourced from the ZIRC Zebrafish International Resource Center (https://zebrafish.org).

### Plasmids and cloning

Human EWSR1::FLI1 coding sequence was provided by Chris Denny, University of California-Los Angeles, USA ^63^. The Gateway expression system (Invitrogen) was used for generation of all constructs for expression in zebrafish ^64^. GFP-2A-EWSR1::FLI1 flanked by attB2r site (at 5′ primer) and attB3 site (at 3′ primer) was cloned into a 3′ entry vector according to the provided protocol ^65^. The plasmids containing a STOP-DsRed-STOP sequence were a generous gift from Eric Olson. Likewise, DsRed-STOP-GFP-2A-EWSR1::FLI1 coding sequence flanked by attB1 and attB2 sites was cloned into a middle entry vector ^64^. The *ubi* promoter was a kind gift from Len Zon (Addgene #27320) ^66^. The final ubi-DsRed-stop-GFP-2A-EWSR1::FLI1 construct was previously described and characterized ^16^. The destination vector pDestTol2pA2 and 3′ SV40 late poly A signal construct were used for construct generation by an LR reaction with LR Clonase II Plus (Invitrogen) ^64^. Transposase and Cre RNAs were synthesized from plasmids pCS2FA and pCS2-Cre.zf using the mMessage mMachine kit (Applied Biosystems/Ambion, Foster City, CA).

The -4.9 sox10 enhancer was cloned into a 5’ entry vector and used to generate -4.9 sox10:mTagBFP-polyA and -4.9 sox10:Cre plasmids ^20^.

### Zebrafish microinjection

Zebrafish embryos were injected at the single-cell stage. The injection mixes contained 50 ng/µl of Tol2 transposase mRNA, 10–50 ng/µl of described DNA constructs, 0.1% phenol red, and 0.3× Danieau’s buffer. The injection mixes containing ubi:RSG-2A-EWSR1::FLI1 also contained 0.5 ng/µl of *Cre* RNA.

### Morpholino injection

Morpholinos were injected in one-cell embryos. The sequence and amount of antisense morpholino oligonucleotides (Gene Tools) used in this study were as follows: *tfap2a*-MO (5 ng): 5’-GCGCCATTGCTTTGCAAGAATTG-3’; combination of two *foxd3* morpholinos *foxd3*- MO^5′UTR^ (1.5 ng): 5’- CACCGCGCACTTTGCTGCTGGAGCA -3’ and *foxd3*-MO^AUG^ (1.5 ng): 5’- CACTGGTGCCTCCAGACAGGGTCAT -3’ ^22,60^. Standard Control Oligos were used as control (Gene tool). The combination of three morpholinos or Standard Control Oligos were co-injected alone, or with ubi:RSG-2A-EF1 and *Cre* mRNA.

### Zebrafish tumor collection and processing for histology

Zebrafish were examined under a Leica fluorescent stereomicroscope to identify the presence of GFP-positive tumors. Zebrafish that exhibited no GFP fluorescence were classified as not showing EWSR1::FLI1-dependent tumor formation. Fish bearing tumors were euthanized and fixed in a 4% paraformaldehyde/1× phosphate-buffered saline (PBS) for 48 hours at 4°C. Subsequently, the fish underwent a 5-day decalcification process in 0.5M EDTA. Zebrafish were then processed and mounted in paraffin blocks for sectioning and further experiments.

### Immunohistochemical staining

Slides with paraffin-embedded tissue sections were baked for 60 min at 60°C, immersed with xylene, re-hydrated with 100% ethanol, 95% ethanol, 75% ethanol and distilled H_2_O two times each for 5 min each. Antigen retrieval was performed in Trilogy reagent (920 P, Sigma) for 10 min in a pressure cooker. Slides were cooled and blocked with 3% H_2_O_2_ for 30 min, followed by blocking with 1%BSA/1× PBST for 1 hr. Slides were incubated with primary antibodies Anti- CD99 antibody (ab108297, Abcam) at 1:200 and Anti-FLI1 (ab15289, Abcam) at 1:100 overnight. Secondary antibodies were Anti-rabbit IgG, HRP-linked Antibody (7074 S, Cell Signaling) and Anti-mouse IgG, HRP-linked Antibody (7076 S, Cell Signaling). SignalStain DAB Substrate Kit #8059 was used for chromogen staining according to the manufacturer’s instructions. Slides were also stained with hematoxylin and eosin or Periodic acid–Schiff stain, dehydrated, and mounted with Permount mounting media. The staining was repeated more than three times.

### RNAscope whole mount in situ hybridization

For in situ hybridizations, we employed the RNAscope approach ^67^. Zebrafish embryos were fixed using 4% PFA at 24 hpf, 72 hpf, 7dpf, or 14dpf and were subsequently utilized for a RNAscope Fluorescent Multiplex V2 Assay according to the manufacturer’s ‘RNAscope assay on Whole Zebrafish embryos’ protocol with slight adaptations. All hybridization stages took place in a 40°C water bath. During each washing step, samples were gently agitated using 1 ml of 1× Wash Buffer, with two washes performed for each step for 5 minutes each. The following probes were used for this study: Hs-EWSR1-FLI1-No-XDr-C1 for specific detection of human EWSR1::FLI1, EGFP- C1, Dr-ta-C1, Dr-nkx2.2a-C2, Dr-ta-C2, Dr-hoxd13a-C2, Dr-hoxa13b-C2, Dr-isl2b-C2, Dr- neurog1-C2, Dr-elavl4-C2, Dr-tbx5a-C3, Dr-fgf8a-C3, Dr-sox10-C3, Dr-fgf10b-C3, Dr-fgf10a- C4, Dr-eomesa-C4, Dr-tbx4-C4, Hs-EWSR1-FLI1-No-XDr-C4.

### Imaging

All imaging was performed on Leica M205 FA stereomicroscope, Leica Thunder, Leica S8APO stereomicroscope, Leica DM4000B, or Leica STELLARIS 5 using LAS X software. Embryos were imaged at 12 hpf, 24 hpf, 48 hpf, 72 hpf, and 5 dpf using a Leica fluorescent stereomicroscope. Time-lapse movies were created using a Leica M205 FA stereomicroscope equipped for epifluorescence, starting from 6-8 up to 24 hpf and from 30 to 72 hpf. Images of whole-mount embryos stained with RNAscope approach were captured on a Leica STELLARIS 5 confocal microscope, and slides were imaged on a Leica DM4000B.

### scRNAseq and snATACseq library preparation and alignment

Trunks from converted -*28.5Sox10:Cre; actab2:loxP-BFP-STOP-loxP-DsRed (Sox10>DsRed)* or -28.5*Sox10:Cre;actab2:loxP-BFP-STOP-loxP-DsRed; Ubi:loxP-DsRed-STOP-loxP-GFP-2A- EF1 (Sox10>GFP-2A-EF1)* fish at 7 dpf were cut posterior to the ear and excluded swim bladder at all stages, further cut into multiple pieces, and 24 trunks were placed per tube in cold 1x PBS. Tumors from *Sox10>GFP-2A-EF1* fish or normal tissue from control fish were dissected at young adult stages, cut into multiple pieces, and placed in cold 1x PBS. 1x PBS was then removed and replaced with pre-warmed Accumax Cell Aggregate Dissociation Medium. Samples were placed on a nutator at 30-32 °C, followed by mechanical dissociation by pipetting every 10 min (Innovative Cell Technologies Inc, AM105) for 1.5-2 hr. Cells were pelleted (200 × *g*, 10 min, 4 °C), and supernatant was removed (leaving roughly 200 µL) and replaced with 1 mL of cold Hank’s Buffer solution (0.25% BSA, 10mM HEPES, 1x HBSS). Cells were vortexed to resuspend and pelleted again (376 × *g*, 5 min, 4 °C), and supernatant was removed (leaving roughly 100 µL) and replaced with 100 µL of cold 1x PBS per tube . Pellets were resuspended by vortexing and filtered through 40 µm strainers (Corning, 431750). Cells were then resuspended, placed on ice, and subjected to fluorescence-activated cell sorted (FACS). FACS sorting was performed on a BD FACSymphony S6 6-way cell sorter for only DsRed+ or DsRed+/BFP+ for control animals and GFP+ or GFP+/DsRed+ for EWSR1::FLI1+ animals (Table S1).

For snATACseq library construction, sorted cells were subjected to nuclei isolation according to manufacturer’s instructions (10x Genomics, protocol CG000169, “Low Cell Input Nuclei Isolation”), followed by integrity check of DAPI-stained nuclei under a confocal microscope (40x objective) before library synthesis. Barcoded single-nuclei ATAC libraries were synthesized using 10x Genomics Chromium Single Cell ATAC Reagent Kit v1.1 per manufacturer’s instructions. Libraries were sequenced on Illumina NextSeq (500/550 Mid Output v2.5 (150 cycles)), or HiSeq (3000/4000 SBS Kit (150 cycles), FC-410-1002 + 3000/4000 PE Cluster Kit, PE-410-1001) machines at a depth of at least 100,000 reads per nucleus for each library. Both read1 and read2 were extended from 50 cycles, per the manufacturer’s instruction, to 65 cycles for higher coverage. Cellranger ARC v2.0.0 (10x Genomics) was used for alignment against GRCz11 (built with GRCz11.fa, JASPAR2020, and GRCz11.105.gtf), peak calling, and peak-by-cell count matrix generation with default parameters.

For multiome library construction, FACS-sorted cells from EWSR1::FLI1+ and control fish at 7 dpf and tumor and control tissue from young adult fish were subjected to nuclei isolation protocol performed per manufacturer’s instructions (10x Genomics, protocol CG000365 Rev A, “Low Cell Input Nuclei Isolation”) followed by integrity check of DAPI-stained nuclei under a confocal microscope (40x objective) before library synthesis. Barcoded single-nuclei multiome (ATAC and cDNA) libraries were synthesized using 10x Genomics Chromium Next GEM Single Cell Multiome ATAC + Gene Expression (PN-1000285). Libraries were sequenced on Illumina NextSeq or HiSeq machines at a depth of at least 100,000 reads per nucleus for each library. When sequencing the multiome ATAC library from EWSR1::FLI1+ and control cells (7 dpf), Read4 was extended from 100 cycles, per the manufacturer’s instruction, to 102 cycles for higher coverage. When sequencing the multiome GEX (cDNA) library from EWSR1::FLI1+ and control cells (7 dpf), Read1 and Read 4 were both extended from 58 cycles, per the manufacturer’s instruction, to 59 cycles for higher coverage. Cell Ranger ARC v2.0.0 and Cell Ranger v8.0 (10x Genomics) were used with 7 dpf and adult multiome datasets, respectively, for alignment against GRCz11 and a customized version of GRCz11 that includes eGFP-2A-EF1 (built with GRCz11.fa and GRCz11.105.gtf) and gene-by-cell count matrices were generated with default parameters. Cellranger RNA v6.0.2 (10x Genomics) was used to generate snRNA-only datasets from the multiome experiments (must indicate --chemistry=ARC-v1 and --include-introns). Cellranger ARC v2.0.0 (10x Genomics) was used to generate snATAC-only datasets from the multiome experiments (must indicate --chemistry=ARC-v1) (Table S1).

### Data processing of scRNAseq and snATACseq

The count matrices of both multiome and snATACseq data were analyzed by R (v4.1.3) package Seurat (v4.3.0) and Signac (v1.6.0). For snRNAseq (Fig. 4), 7 dpf EWSR1::FLI1+ and 7 dpf control trunk datasets were merged (merge, Seurat-methods), matrices were normalized (NormalizeData) and scaled for the top 2000 variable genes (FindVariableFeatures, method = “vst” and ScaleData). The scaled matrices were dimensionally reduced to 20 principal components (RunPCA) based on JackStrawPlot and ElbowPlot, in addition to prior biological knowledge of the neural crest contribution to the trunk ^68^. The data were then subjected to neighbor finding (FindNeighbors) and clustering (FindClusters, resolution = 0.4). The data were visualized through UMAP with the first through 20th principal components as input. For snATACseq datasets (Fig. 5), Cellranger (10x Genomics) 7 dpf EWSR::FLI11+ and 7 dpf control trunk aggregated matrices were normalized using RunTFIDF and RunSVD functions. The neighbor finding, clustering, and visualization were performed (RunUMAP, FindNeighbors, FindClusters, algorithm = 3 for FindClusters with resolution = 0.8) with input of the second to thirtieth LSIs. To test the enriched genes, pseudo-gene activities, and/or accessible chromatin regions in both snRNAseq and snATACseq data, likelihood-ratio tests were performed through the FindAllMarkers function (min.pct = 0.2, test.use = ‘bimod’, log.fc =0.03) ^69^ with cutoff of adjusted *p* value smaller than 0.05. For snRNAseq (Fig. 3), tumor and control datasets were merged (merge, Seurat-methods), matrices were normalized (NormalizeData) and scaled for the top 500 variable genes (FindVariableFeatures, method = “vst” and ScaleData). The scaled matrices were dimensionally reduced to 15 principal components (RunPCA) based on ElbowPlot.

### CUT&RUN

Embryos at 24hpf or tumors positive for GFP-2A-EF1 were dissected, cut into multiple pieces, and placed in pre-warmed Accumax at 30-32°C, followed by mechanical dissociation by pipetting every 10 min (Innovative Cell Technologies Inc, AM105) for 1-1.5 hr. Cells were pelleted (200 × *g*, 10 min, 4 °C), and supernatant was removed and replaced with 1 mL of cold Hank’s Buffer solution (0.25% BSA, 10mM HEPES, 1x HBSS). Cells were resuspended by pipetting up and down and pelleted again (376 × *g*, 5 min, 4 °C), and supernatant was removed and replaced with 1ml of 1x PBS. Pellets were resuspended and filtered through 40 um strainers (Corning, 431750). Cell number was assessed by automated cell counter. Cells were fixed in 1%PFA for 2 min. 100,000-200,000 cells per sample were aliquoted and pelleted for processing using CUT&RUN Assay Kit #86652 according to the manufacturer’s protocol.

Sequencing read quality was examined using FastQC. Trimming of low-quality reads and clipping of sequencing adapters were done using the FASTQ Quality Trimmer (Galaxy Version 1.1.5). Reads were aligned to the zebrafish genome (GRCz11) using Bowtie2 (Galaxy Version 2.5.0) using default settings. Bam files were filtered with Filter SAM or BAM (Galaxy Version 1.8) and sorted SortSam (Galaxy Version 2.18.2.1). Duplicate reads were removed using RmDup (Galaxy Version 2.0.1). The individual bam files corresponding to biological replicates were merged using the MergeSamFiles (Galaxy Version 2.18.2.1) command. CUT&RUN peaks were called using MACS2 (Galaxy Version 2.2.7.1) with the callpeak command (--nomodel, --extsize 150, --shift 0) and a q-value threshold of <1e-05 for all Input_embryos, FLAG_embryos, Input_tumor, and FLAG_tumor files.

### Motif enrichment analysis and peak annotation

Open chromatin regions were determined from the snATACseq datasets using AccessiblePeaks (Signac, min.cells = 10). Common peaks between the 7 dpf EWSR1-FLI1+ and 7 dpf control trunk datasets were determined using bedtools intersect function on all accessible peaks (Signac, AccessiblePeaks). Differentially accessible peaks were identified from a merged dataset in Loupe Browser (10x Genomics) as “globally distinguishing” peaks between samples (by Library ID), with all peaks p-value > 0.05 removed. HOMER de novo motif enrichment ^70^ was then performed using findMotifsGenome.pl (-size given) with motif length set to default (-motif length 8,10,12) and the common peak file set as background (-bg) (Fig. 5C). HOMER de novo motif enrichment analysis on accessible peaks from Tumor snATACseq dataset was then performed using findMotifsGenome.pl (-size 200) and the control peak file set as background (-bg) (Fig. 5E). For CUT&RUN data analysis, identified peaks from MACS2 (Galaxy Version 2.2.9.1) were used for HOMER de novo motif analysis using findMotifsGenome.pl (-size given -mask) and the input peak file set as background (-bg) (Fig. 5F). The annotation of ETS sites was performed by HOMER using the annotatePeaks.pl function. Open chromatin regions were determined from the snATACseq datasets using AccessiblePeaks (Signac, min.cells = 10). Peaks specific to 7 dpf control dataset compared to 7 dpf EWSR1::FLI1+ were identified using bedtools intersect (-v), and vice versa for peaks specific to the 7 dpf EWSR1::FLI1+ dataset. Then, annotatePeaks.pl was run using an ETS motif file (NRYTTCCTGH) using seq2profile.pl. Number of motif instances per peak was calculated and plotted within Microsoft Excel (Fig.5D).

### Statistics

Statistical analysis was performed using GraphPad Prism 9 (La Jolla, CA). The number of individual experiments, replicas, and samples analyzed, and significance is reported in the figure legends. Statistical significance was calculated by Student’s *t*-test for two-group comparison, one-way analysis of variance for comparison of multiple groups with one control group and for comparison between different experimental groups. p > 0.05 = n.s., *p < 0.05, **p < 0.01, ***p < 0.001, and ****p < 0.0001.

## Acknowledgments

We thank Hillary Mahon, Megan Matsutani and Maya Lujan for fish care, and Anna Luzzi for laboratory management. We thank the Children’s Hospital Los Angeles (CHLA) Translational Pathology Core, supported by the USC Norris Comprehensive Cancer Center grant P30CA014089 from the National Institutes of Health (NIH), and the Cellular Imaging and Spatial Biology and Genomics Cores of The Saban Research Institute (TSRI) at CHLA for exceptional services and for their expertise. We also thank Rosa Uribe for sharing a protocol of tissue dissociation. E.V. also thanks Vladimir Zhemkov for support and discussions. E.V. was supported by a K99/R00 Pathway to Independence Award 1K99CA270282 from NIH NCI, by a Young Investigator Grant from Alex’s Lemonade Stand Foundation (Grant 22-27160), and by a Research Career Development Fellowship from TSRI. J.F.A. was supported by grants P30CA014089 and U54CA231649 from NIH, by the 1Million4Anna and Curing Kids Cancer Foundations, and by the Dr. Kenneth O. Williams Chair in Cancer Research at Children’s Hospital Los Angeles.

## Author Information

Contributions

E.V., C.A., J.G.C. and J.F.A. conceived the experiments. E.V., C.A., Y.L. and R.B. carried out experiments. E.V. and C.A. performed data analysis. E.V. and J.F.A. obtained funding. J.G.C. and J.F.A. oversaw the project. E.V., J.G.C., and J.F.A. wrote the manuscript. All authors edited and approved the manuscript.

Corresponding Authors

Correspondence and requests for materials should be addressed to J. Gage Crump and James F. Amatruda.

## Ethics Declarations

Competing Interests

The authors declare no competing interests.

## Supplementary Materials

Table S1: Overview of Single Cell Transcriptome and Multiome parameters

Fig.S1 Fig.S2 Fig.S3 Fig.S4 Fig.S5 Fig.S6

## Supplemental Movies

**Movie 1 -** Time-lapse imaging of embryos mosaically expressing GFP-2A or GFP-2A-EF1, starting from 5 hpf (approximately 50% epiboly stage) to 24 hpf.

**Movie 2** – 3D analysis of z-stack confocal images of larvae shows patterns of mosaic Cre-inducible GFP (control) or GFP-2A-EF1 at 24 hpf.

**Movie 3** - Time-lapse imaging of embryos mosaically co-expressing GFP-2A-EF1 and - 4.9Sox10:mTagBFP2 starting from 10 hpf to 24 hpf.

**Movie 4** - 3D analysis of z-stack confocal images of larvae with outgrowth at 14dpf subjected to double RNAscope staining for *tbxta (red)* and *neurog1* (white).

**Movie 5** – 3D analysis of z-stack confocal images of larva with outgrowth at 72 hpf, subjected to double RNAscope staining for *neurog1* (red) and *EWSR1::FLI1* (green).

Table S1:

**Figure S1.**
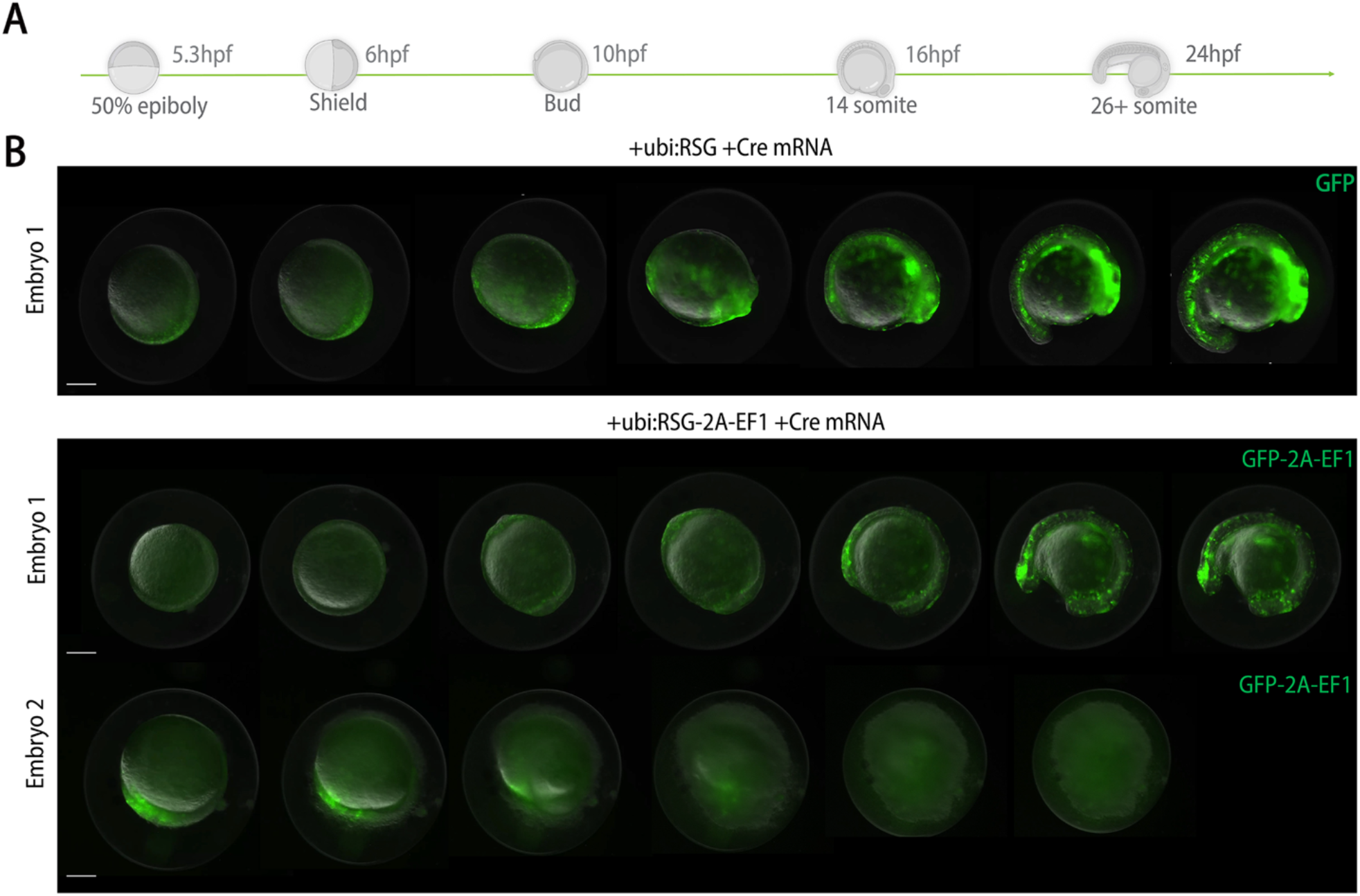
(A) Schematic of zebrafish development during the first 24 hpf. **(B)** Serial imaging of embryos mosaically expressing GFP (top panel) or GFP-2A-EF1, starting from 5 hpf (approximately 50% epiboly stage) up to 24 hpf. Scale bars, 200 μm.

**Figure S2.**
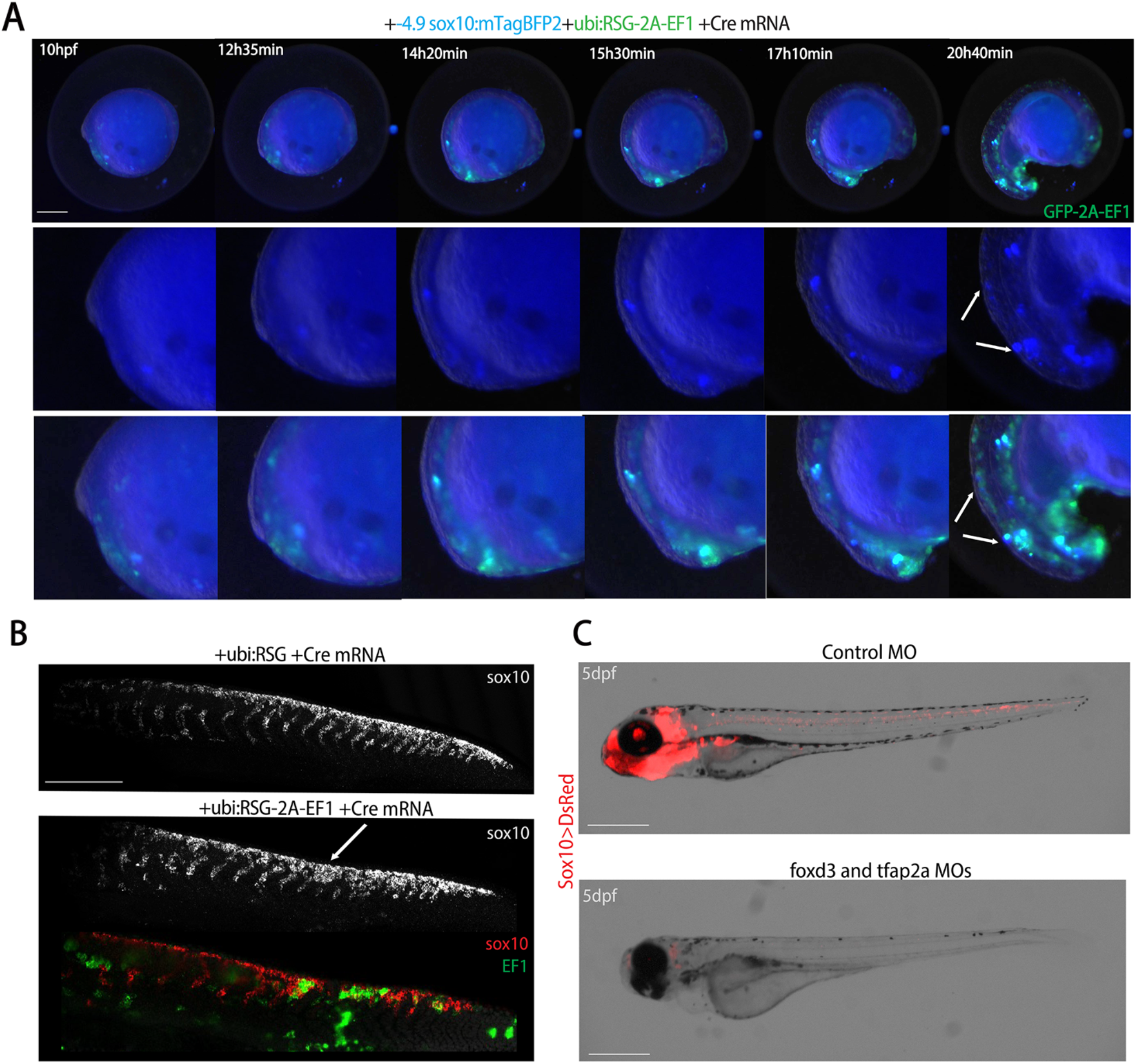
(A) Serial imaging of embryos mosaically co-expressing GFP-2A-EF1 and - 4.9sox10:mTagBFP2 from 10 to 24 hpf. Arrows denote double-positive cells. Scale bars, 200 μm. **(B)** RNAscope in situ hybridization for *sox10* and *EWSR1::FLI1* in embryos mosaically expressing GFP (control, top panel) or GFP-2A-EF1 (bottom panel). Arrow indicates an example of *sox10* and *EWSR1::FLI1* co-localization. Scale bars, 200 μm **(C)** Validation of NCC loss by injecting control (top panel) or *foxd3*/t*fap2a* morpholinos (bottom panel) into Sox10>DsRed transgenic fish Scale bars, 500 μm.

**Figure S3.**
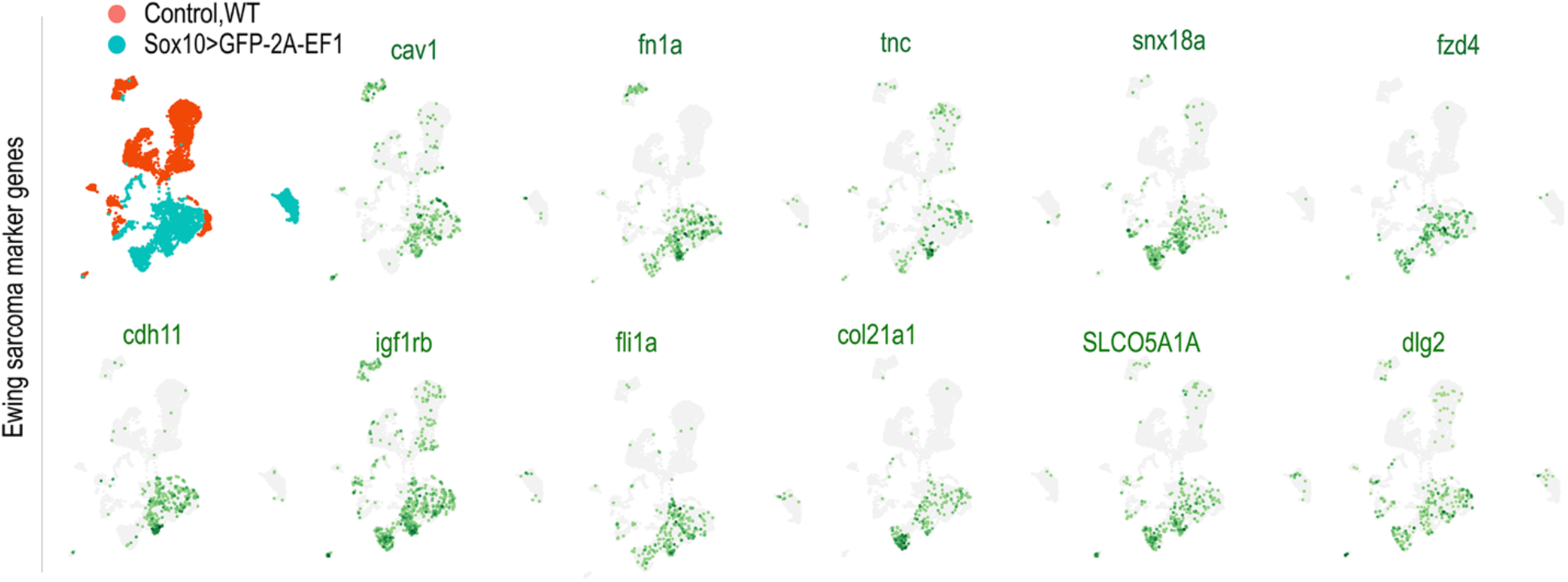
Feature plots show expression of the zebrafish orthologs of human Ewing sarcoma markers in combined control and EWSR1::FLI1+ adult tumors from the NCC-specific model.

**Figure S4.**
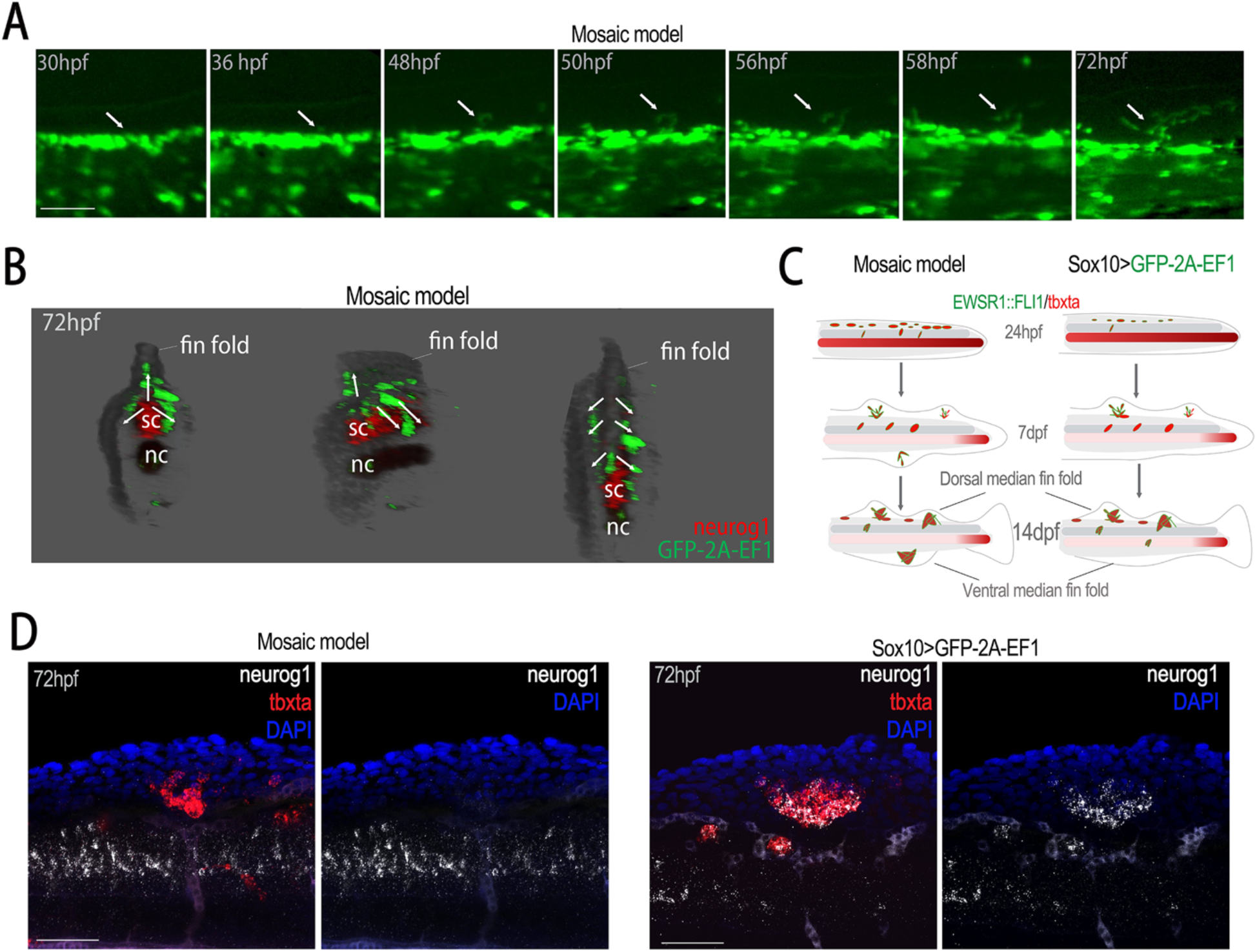
(A) Time-lapse imaging of the fin fold region of an embryo mosaically expressing GFP-2A-EF1 from 30 to 72 hpf. Arrows indicate the outgrowth. Scale bars, 50 μm. **(B)** In situ hybridization for *neurog1 (red)* and *EWSR1::FLI1 (green)* in a 72 hpf larva with mosaic misexpression of GFP-2A-EF1. As opposed to the NCC-specific model, *neurog1* is not co-expressed with the oncofusion in the mosaic model. **(C)** A schematic showing differences between mosaic and Sox10>GFP-2A-EF1 models. **(D)** In situ hybridization for *neurog1 (white)* and *tbxta* (red) show co-localization in the dorsal fin fold masses in the NCC-specific model (Sox10>GFP- 2A-EF1) but not the mosaic model. Scale bars, 50 μm.

**Figure S5.**
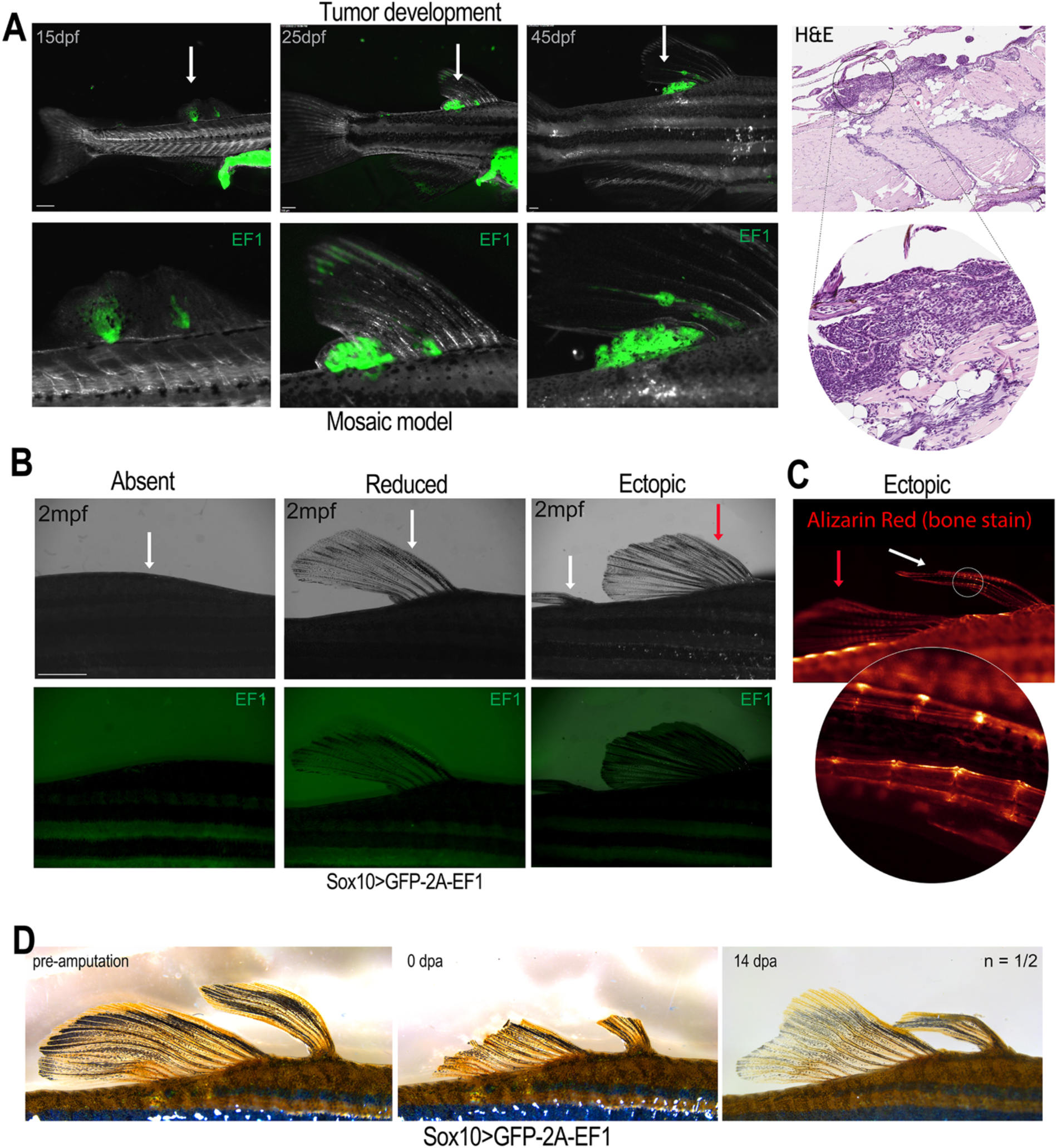
(A) Representative serial images of a GFP-2A-EF1-positive outgrowth that transforms into a tumor from 15 to 45 dpf. H&E staining of tumor sections from the same fish. Scale bars, 300 μm. **(B)** Images of fish with absent fin (white arrow), reduced fin (white arrow), and multiple (white arrow indicates ectopic fin, red arrow indicates normal fin) in Sox10>GFP-2a-EF1 fish. Green channel shows absence of GFP-2A-EF1expression. Scale bars, 2 mm. **(C)** Alizarin Red staining of ectopic fin in Sox10>GFP-2A-EF1 fish, with magnified region highlighting presence of normal joints in the fin rays. (white arrow indicates ectopic fin, red arrow indicates normal fin). **(D)** Regeneration of EWSR1::FLI1-induced ectopic fins by 14 days post-amputation (dpa).

**Figure S6.**
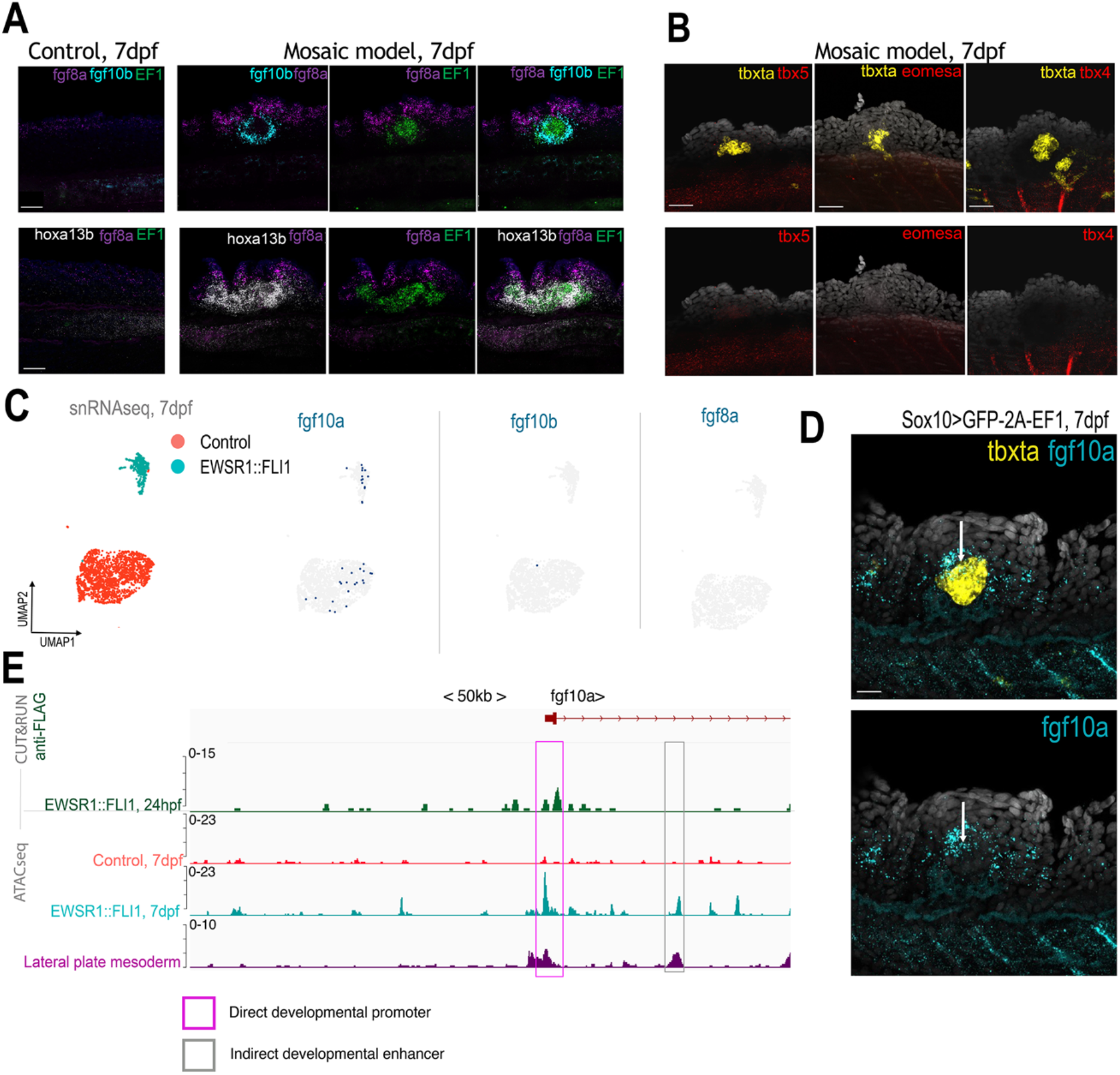
(A) Expression of *fgf8a*, *fgf10b*, *EF1*, and *hoxa13b* in dorsal fin fold of control and mosaic model (ubi:RSG-2A-EF1+Cre mRNA) larvae. Scale bars, 40 μm. **(B)** *tbxta* but not *tbx5*, *eomesa*, or *tbx4* is expressed in outgrowths in larvae of the mosaic model. Scale bars, 40 μm. **(C)** Feature plots show expression of *fgf10a* but not *fgf10b* and *fgf8a* in combined control and EWSR1::FLI1+ datasets at 7 dpf . **(D)** In situ hybridization shows that expression of *fgf10a* partially overlaps with *tbxta* in dorsal fin fold outgrowth of the NCC-specific model at 7 dpf. Arrow denotes region of overlap. Scale bars, 20 μm. **(E)** Integrated coverage plots for CUT&RUNseq (EWSR1::FLI1+ at 24 hpf) and snATACseq (control and EWSR1:: FLI1 at 7 dpf, and lateral plate mesoderm downloaded from GEO Series GSE243256). Genomic locus for *fgf10a* is shown.

